# Evidence for a general neural signature of face familiarity

**DOI:** 10.1101/2021.04.18.440317

**Authors:** Alexia Dalski, Gyula Kovács, Géza Gergely Ambrus

## Abstract

We explored the neural signatures of face familiarity using cross-participant and cross-experiment decoding of event-related potentials, evoked by unknown and experimentally familiarized faces from a set of experiments with different participants, stimuli, and familiarization-types. Participants were either familiarized perceptually, via media exposure, or by personal interaction. We observed significant cross-experiment familiarity decoding involving all three experiments, predominantly over posterior and central regions of the right hemisphere in the 270 – 630 ms time window. This shared face familiarity effect was most prominent between the Media and Personal, as well as between the Perceptual and Personal experiments. Cross-experiment decodability makes this signal a strong candidate for a general neural indicator of face familiarity, independent of familiarization methods and stimuli. Furthermore, the sustained pattern of temporal generalization suggests that it reflects a single automatic processing cascade that is maintained over time.

**Highlights:**

- We investigated if a general neural signature of face familiarity exist
- A cross-experiment decoding analysis of EEG data was used
- The analysis involved perceptual, media and personal familiarization methods
- We found a preserved pattern of familiarity decoding across experiments between 270 and 630 ms post-stimulus
- This signature is consistent with previous reports on face familiarity effects

## Introduction

The successful recognition of a face requires the interplay of multiple cognitive systems via connections across several parts of the brain. A number of regions have been shown to highly favor face stimuli, while others are known to carry out more general-purpose processing (Duchaine and Yovel, 2015). The joint, synchronized functioning of this network enables the integration of perceptual, episodic and recognition memory, semantic, contextual and affective information that underlies everyday interactions with others (Kovács, 2020).

We know relatively little about how this system accommodates the addition of a new face-identity to the existing repertoire of previously learned faces. To study the process of becoming familiar with someone, it is a good starting point to ask if a neural signature exists that reliably flags unknown and/or known faces, irrespective of the mode of acquisition or depth of encoding. Furthermore, should such signals exist, are they shared across individuals, implying more of an automatic processing cascade, or do they rather arise as a combination of stimulus properties and idiosyncratic activation patterns, shaped by the person’s prior experience and functional-anatomical mappings (Wilmer et al., 2010; Verhallen et al., 2016; Sanchez et al., 2021)?

While the answer is most certainly a combination of both, compelling evidence exists for the markedly differential processing of familiar and unfamiliar faces (for a review see e.g. (Johnston and Edmonds, 2009)). For example, in the macaque brain, images of personally familiar conspecifics engage the face processing network more than unfamiliar ones, and recruit additional areas in the perirhinal cortex and temporal pole that rapidly activate when familiar faces became recognizable (Landi and Freiwald, 2017). Strong evidence from patients with acquired prosopagnosia (a brain-lesion-caused inability to recognize or experience a conscious feeling of familiarity with the faces of people previously encountered) provide clues that face familiarity can be inferred from physiological signals even without conscious recognition. Tranel and Damasio (1985) describe two female patients with bilateral occipito-temporal lesions who have shown elevated electrodermal responses for personally familiar faces even in the absence of recall. Conversely, in Capgras syndrome person identification remains preserved but the associated feeling of familiarity appears to be lost and, in contrast to prosopagnosia, skin conductance responses do not differentiate between familiar and unfamiliar faces (Ellis and Lewis, 2001). In healthy volunteers, eye movement patterns reveal distinct markers of familiar face recognition (Rosenzweig and Bonneh, 2019) that may even defy active countermeasures at concealment (Millen and Hancock, 2019).

In a recent study, Yan, Young and Andrews (2017) demonstrated that while the processing of unfamiliar faces is not entirely automatic, familiarity with a face does increase the automaticity for certain facial characteristics. In a matching task, participants were instructed to report whether certain properties (sex, identity, ethnicity, and expression) were the same or different for two briefly presented faces. Automaticity was tested by comparing performance between cases when the instruction was given at the beginning of the trial to when it was given at the end of the trial. While performance for ethnicity (white/black) and expressed emotion (neutral/happy) benefited from cues at the beginning of the trial for both familiar and unfamiliar faces, no differences were found for judgments of sex and identity from familiar faces (celebrities), indicating a strong automatic processing of these dimensions. In agreement with this finding, Dobs et al. (2019) observed that familiarity enhanced the decodability of sex and identity (but not age) information at early stages of visual processing, suggesting that this familiarity enhancement effect is due to these processes being tuned to familiar face features. In a change-detection task, that did not require explicit image recognition, Buttle and Raymond (2003) presented briefly and successively pairs of faces (either famous or recently learned), changing one of the two faces between the presentations, and found that the change-detection performance was significantly better for famous when compared to unfamous faces. Also, the authors found a clear left visual filed bias, consistent with the view that the right hemisphere is involved in a more global mode of face processing, whereas the left hemisphere attends to local information, suggesting that extensive experience with a face gives rise to a more configural as opposed to a featural mode of face processing. Measuring visual search performance with familiar (male Hollywood actors) and unfamiliar faces in paradigm, Persike et al. (2013) found that face familiarity also strongly enhances performance, both in terms of accuracy and reaction time. While for target-absent trials only a small familiarity advantage was shown, for target-present trials the effect was pronounced. Moreover, participants needed more than twice the time to process one item of unfamiliar, compared to familiar faces. The authors concluded that familiarity enhances difference detection among faces via efficient global mechanisms, connected to long term memory representations.

As we have seen, compelling evidence exists for differential processing of familiar and unfamiliar faces, implying the existence of automatic processes that provide prioritized access to certain stimulus characteristics when compared to faces of unknown individuals. The next logical question is, does this processing happen in a cascade-like manner with each step pre-wired, in temporal order, linked to the activity of distinct areas of the brain?

If signals indicative of an automatic processing cascade exist, these should be similar across individuals, and also robust to the properties of the stimuli, as well as to the type of familiarization. As an extreme, and somewhat simplified example, we should find these hypothesized neural signatures in Alice when she meets her grandmother for a coffee, and in Bob, when he sees a known celebrity on the cover of a magazine while causally browsing at a newsstand.

Of the various neuroimaging methods, magnetoencephalography (MEG) and electroencephalography (EEG) provide the adequate temporal resolution to uncover the temporal dynamics of face-familiarity processing in the human brain. Previous research has identified several electrophysiological components that appear to be indicative of the processing of face stimuli. The earliest of these components, the P1 (around 70 - 110 ms, with typically occipital peak) and the N170 (around 160 - 200 ms, with a lateral-parieto-occipital peak) most probably reflect early perceptual stages as they appear to be strongly modulated by the physical properties of the stimuli (Ambrus et al., 2019b). It is important to note that familiarity information may indeed already be present in this time-window, but due to methodological constrains, its signals do not reach the surface sensors, or they are drowned out by the strong response produced by the rapid processing of physical features immediately after stimulus onset. A strong indication of this is the fact that in some cases even these early waves have been reported to be influenced by familiarity (Debruille et al., 1998; Caharel et al., 2002; Barragan-Jason et al., 2015), depending on the task, stimulus set, or prior expectations (Johnston et al., 2016). The first ERP component that is behaviorally connected to recognition seems to be the N250, a parieto-occipital negative deflection between ca. 230 - 350 ms. Huang et al. (2017) found a positive correlation between the N250 amplitude and reaction time in a famous/non-famous decision task, suggesting that it is the earliest electrophysiological indicator of familiar face recognition in long-term memory.

More recently, several studies reported finding a late electrophysiological correlate of familiarity in the 400 to 600 ms time range. Using MEG, Dobs et al. (2019) observed a generic familiarity-related component ca. 400 to 600 ms after stimulus onset. Karimi-Rouzbahan et al. (2020) found a similar familiarity-related effect in EEG starting around 200 ms and peaking after approximately 400 ms. Wiese et al. (2019) observed an ERP component (dubbed as sustained familiarity effect, SFE) starting 200 ms after stimulus onset with a maximum between 400 and 600 ms and being the most prominent over bilateral posterior regions for highly familiar versus unfamiliar faces. Finally, in our previous multivariate pattern analysis (MVPA) studies we found image-independent identity effects (Ambrus et al., 2019b) and familiarity effects (Ambrus et al., 2021, in press) falling within the same temporal range.

Conventional univariate ERP analysis techniques contrast evoked response amplitudes between conditions at single or averaged channels, can tell us if the brain processes information differentially at a given time window. Instead of focusing on aggregated responses restricted in space and time, MVPA parses distributed patterns of activity at the trial level. It uses machine learning whereby the evoked responses from trials are iteratively split into training and test sets, and a classifier attempts to classify test trials based on the information it extracted from the training data. The classifier accuracy is therefore indicative of the presence of information in the pattern of evoked responses about the categories of interest (Grootswagers et al., 2017). A further advantage of MVPA over conventional ERP analysis is the flexibility that the train and test data need not necessarily come from the same time point, region of interest, experimental condition, participant, or, as we will see, from the same experiment.

To investigate the shared neural signatures of face familiarity we analyzed ERP data from three different experiments that utilized different (Perceptual, Media, and Personal) familiarization methods. Utilizing a thus far less explored multivariate cross-classification analysis (MVCC, (Kaplan et al., 2015)), we iteratively trained and tested on data from each experiment. This cross-experiment analysis combines elements of cross-modal, cross-participant, and leave-one-participant-out approaches. It is cross-modal in the sense that the mode of familiarization differed across the experiments and cross-participant as the experiments were conducted on different sets of volunteers. It is also similar to a leave-one-participant-out strategy as training is performed on concatenated data from multiple participants and tested on one participant at a time. We believe that this procedure is well-suited to find general neural patterns that are indicative of shared information processing.

## Methods

### Datasets

Three experiments, described in detail in a previous report from our laboratory, form the basis of the present study (Ambrus et al., 2021, in press). These experiments investigated the EEG correlates of face-familiarity in different familiarization conditions, using perceptual, media, and personal familiarization methods.

For details on stimulus presentation, data acquisition and preprocessing, see Ambrus et al. (2021, in press). Briefly, stimuli were ambient face images, in color, depicting originally unfamiliar identities. The images were cropped to center on the inner features of the face, eye aligned, and were presented centrally on a uniform gray background in a pseudorandom order. In all cases, the stimulus presentation time was 600 ms. Spatial and temporal jitter was applied when presenting the stimuli. Participants were given a simple target detection task to ensure maintained attention throughout the sessions. No participant took part in more than one experiment. For the purposes of the current analysis, data for each participant was labelled on the familiarized/unknown dimension.

EEG was recorded using a 64-channel Biosemi Active II system (512 Hz sampling rate; bandwidth: DC 461 to 120 Hz) in a dimly lit, electrically shielded, and sound-attenuated chamber. Data was bandpass filtered between 0.1 and 40 Hz, segmented between −200 and 1300 ms and baseline corrected to the 200 ms preceding the stimulus presentation. The data was downsampled to 100 Hz resolution, and no artifact rejection was performed (Grootswagers et al., 2017). Processing was carried out using MNE-Python (Gramfort et al., 2013).

Below is a short overview of the three experiments described in the report:

1. **Perceptual familiarization**: 42 participants. Stimuli: 4 previously unknown female identities with 10 ambient images each. 40 repetitions of each image. Familiarization: sorting task for two of the 4 identities immediately prior to the EEG recording. The two to-be-familiarized identities were randomly chosen from the 6 possible permutations of the 4 IDs for each participant. Task: detection of slightly rotated images.
2. **Media familiarization**: 24 participants. Stimuli: 4 previously unknown (two male and two female) identities with 2×20 ambient images each, one image repeated 13 times. Familiarization: Watching a TV series with one male and one female person in lead roles between the pre- and post-familiarization EEG recordings. The to-be-familiarized identities were based on the series and were assigned randomly to the participants. Task: detection of repeated images.
3. **Personal familiarization**: 23 participants. Stimuli: 4 previously unknown female identities, with 10 ambient images each, one image repeated 22 times in a session. Familiarization: ca one hour personal interaction with two of the four identities on 3 consecutive days between pre- and post-familiarization EEG recordings. The to-be-familiarized identities were the same for all participants. Task: detection of rotated images.

### Analysis

We used data from the post-familiarization phases of these experiments, using a cross-participant, cross-experiment decoding approach (**Figure 1**). Each iteration of the analysis used data from two out of the three experiments. The epoched EEG from all trials and all channels was concatenated from all participants in one experiment. At each of the 150 time-points (−200 to 1300 ms relative to stimulus presentation onset, sampled at 100 Hz), linear discriminant analysis classifiers (LDA) were trained to categorize familiarized and unknown identities. These classifiers were then used to assess prediction accuracy for familiarity in the other experiment, for each participant separately.

**Figure 1.**
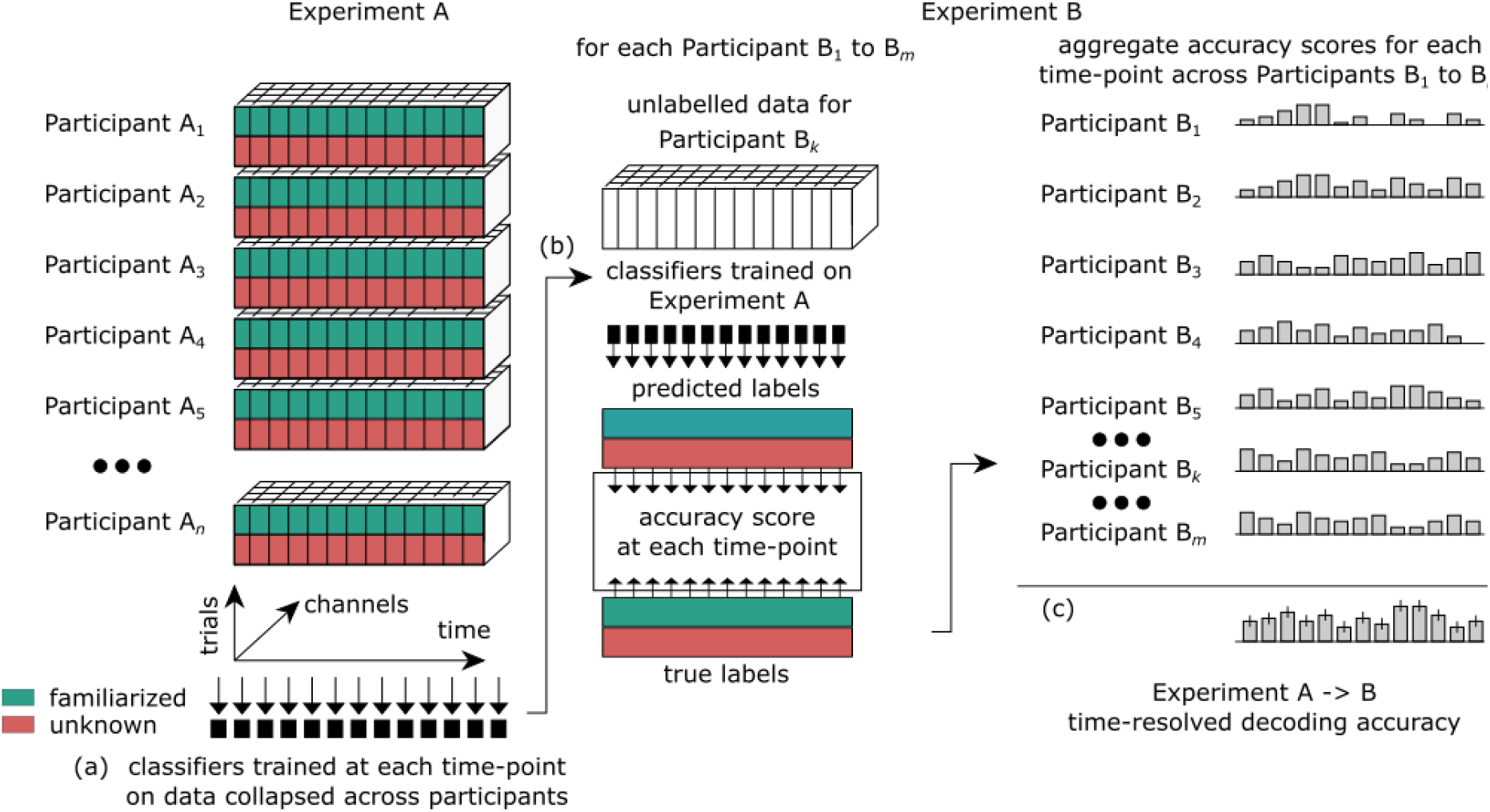
Cross-experiment decoding. (a) Trial × channel × time resolution EEG data from all participants in Experiment A is concatenated. At each time point, a classifier is trained to differentiate between familiarized and unknown trials. (b) These classifiers are then used to categorize the ERPs, at each corresponding time-point in Experiment B, for each participant separately. (c) The subject-level accuracy scores are aggregated to yield the time-resolved decoding accuracy for Experiments A → B.

To investigate the temporal organization of information-processing stages, we used temporal generalization analysis. For this, the classifiers were trained on all timepoints in one experiment and tested on the data at every timepoint in the other, organized in a cross-temporal (training-times × testing-times) decoding accuracy matrix. We reasoned that if a classifier can generalize from one timepoint to another across experiments, similar information-processing may be indicated at those time points (King and Dehaene, 2014).

To gain a finer spatial resolution, we repeated the procedure using a region of interest (ROI) and a searchlight approach. In the ROI analysis, similarly to (Ambrus et al., 2019b) and Ambrus et al. (2021, in press), we pre-defined six scalp locations along the medial (left and right) and coronal (anterior, center, and posterior) planes (see **Supplementary Figure 1**), and used these subsets of channels for training and testing. In the searchlight analysis, all channels were systematically tested by creating quasi-ROIs that also included the neighboring electrodes and conducted the same time-resolved analysis steps as described above (see **Supplementary Figure 2**).

### Statistical testing

In order to increase signal-to-noise ratio, a moving average of 30 ms (3 consecutive time-points) was used on all cross-experiment decoding accuracy data at the participant level (Kaiser et al., 2016; Ambrus et al., 2019b). For the ROI-based analyses, to test if classifiers on a given experiment can decode the data in another experiment, the decoding accuracies were entered into two-tailed, one-sample cluster permutation tests (10,000 iterations) against chance (50%). We also tested cross-experiment decoding accuracies across 6 train/test pairings with two-tailed cluster permutation tests with 10,000 iterations. For the searchlight analysis, two-tailed spatio-temporal cluster permutation tests were used against chance level (50%), with 10,000 iterations.

Additional exploratory analyses were carried out to probe the reliability of decoding on participant-level using a bootstrapping method (Di Nocera and Ferlazzo, 2000; Wiese et al., 2019). In this analysis, predicted labels were randomly reassigned (10,000 iterations) to the trial conditions, and at each reassignment, an accuracy score was calculated. Reliable effects were assumed if a participant’s true accuracy score exceeded 95% of the values, calculated from the random resamplings. Sample-level 95% confidence intervals were calculated the following way: First, *true/false* labels (assuming independent, equal outcomes) were randomly assigned to each participant in the sample, and a sample-wide accuracy score was calculated. This procedure was repeated 50,000 times. We determined the cut-off by ascertaining the number of participants needed to score *true* to reach or exceed the 95% of the accuracy cores in the bootstrapped sample. (n _≥ 95%CI_ = Perceptual: 26 [61.9%], Media: 16 [66.6%], Personal: 15 [65.2%]).

The statistical analyses were conducted using python, MNE-Python (Gramfort et al., 2013), scikit-learn (Pedregosa et al., 2011) and SciPy (Virtanen et al., 2020).

## Results

### All electrodes

Out of the six train-test cross-experiment decoding analyses on the all-electrodes data, significant familiarity decoding effects were observed in three, involving all three experiments, overlapping between 270 and 630 ms (**Figure 2., Figure 5**, black significance markers) and peaking between 330 and 500 ms. Decodability of familiarity was mutual between the Media and the Personal familiarization experiments (Media ↔ Personal), with a significant cluster between 250 and 680 ms, peaking at 380 ms (peak *t*_24_ = 2.86, peak Cohen’s *d* = 0.57) in Personal → Media, and a cluster between 190 and 1010 ms, peaking at 500 ms (peak *t*_22_ = 7.24, peak Cohen’s *d* = 1.51) in Media → Personal direction. Furthermore, in Perceptual → Personal analysis a cluster between 270 and 410 ms with a peak at 330 ms (peak *t*_22_ = 4.36, Cohen’s *d* = 0.91), and between 430 and 630 ms (peak *t*_22_ = 4.11, peak Cohen’s *d* = 0.86) was observed. Although no significant cross-experiment decoding effect was seen between the Media and Perceptual familiarization experiments, for completeness, statistics on these non-significant pairs will be reported as well in the following sections.

**Figure 2.**
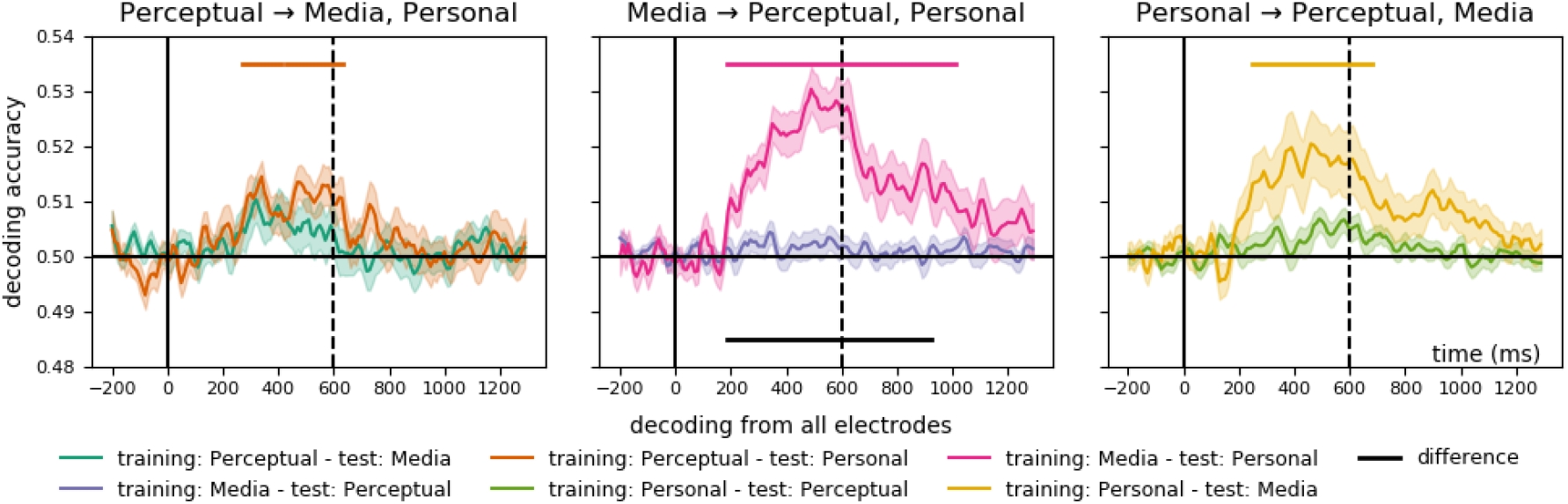
Time-resolved cross-experiment decoding accuracies on all electrodes. Horizontal markers denote clusters with significantly different decoding accuracies against chance (colored markers) and between-pair differences (black markers; two-tailed cluster permutation tests, p < 0.05). Shaded ranges denote standard errors. Note that this analysis is the equivalent of taking the diagonal in the temporal generalization analysis (**Figure 4**.). For detailed statistics on the regions of interest see Supplementary Figure 1 and Supplementary Table 1.

A bootstrapping analysis between the overlapping peak decoding time intervals (320 - 500 ms) revealed that the majority of the participants’ accuracy scores lay above chance level, with 8% (Perceptual → Media), 4.3% (Perceptual → Personal) 47.8% (Media → Personal) and 48% (Personal → Media) of participants’ scores reaching the 95% confidence interval (Figure 3a). To establish an upper bound on subject-level reliability, we repeated the bootstrapping analysis separately for all train/test pairs at their respective peaks. Again, the majority of scores exceeded the chance level in all cases, with 21.4% (Media → Perceptual), 14.3% (Personal → Perceptual), 20% (Perceptual → Media), 34.8% (Perceptual → Personal) 74% (Media → Personal) and 52% (Personal → Media) of participants’ scores over the 95% confidence interval (Figure 3, b).

**Figure 3.**
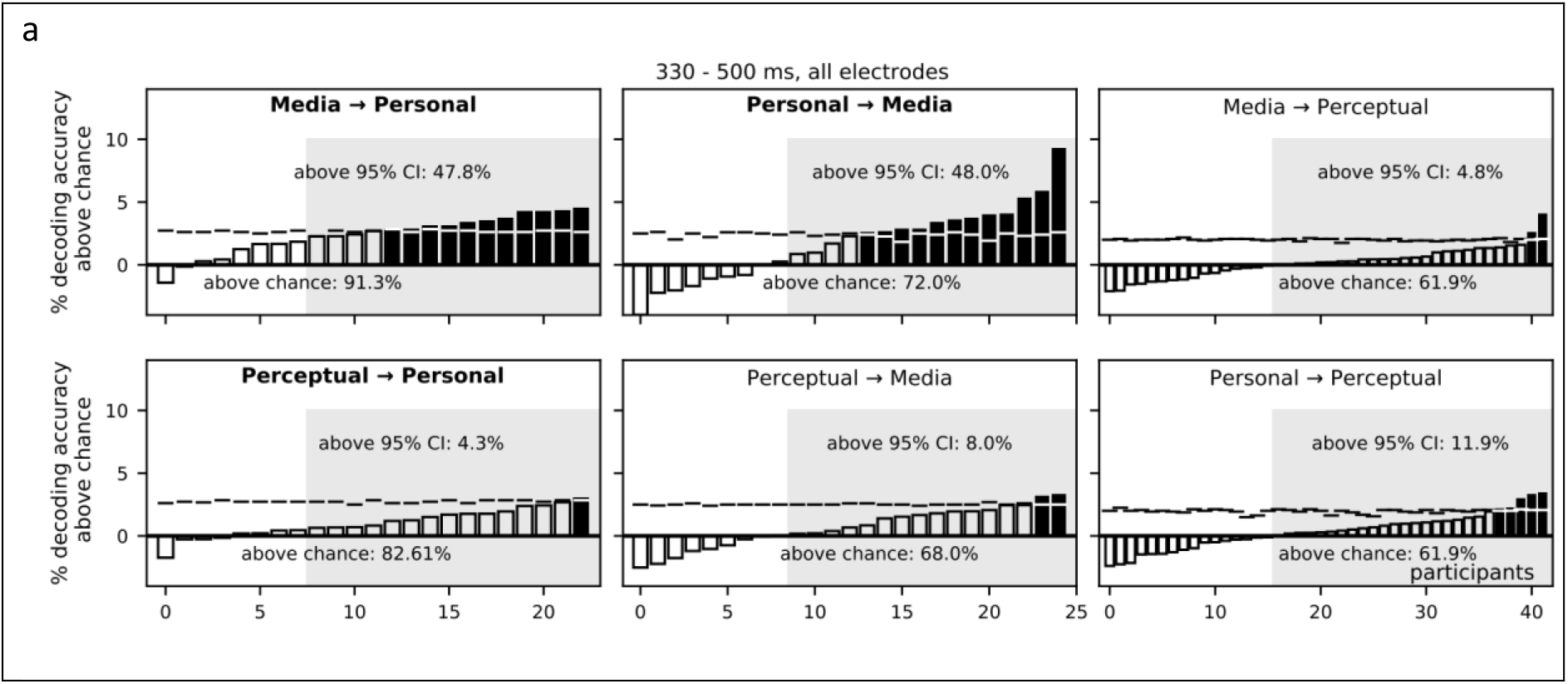
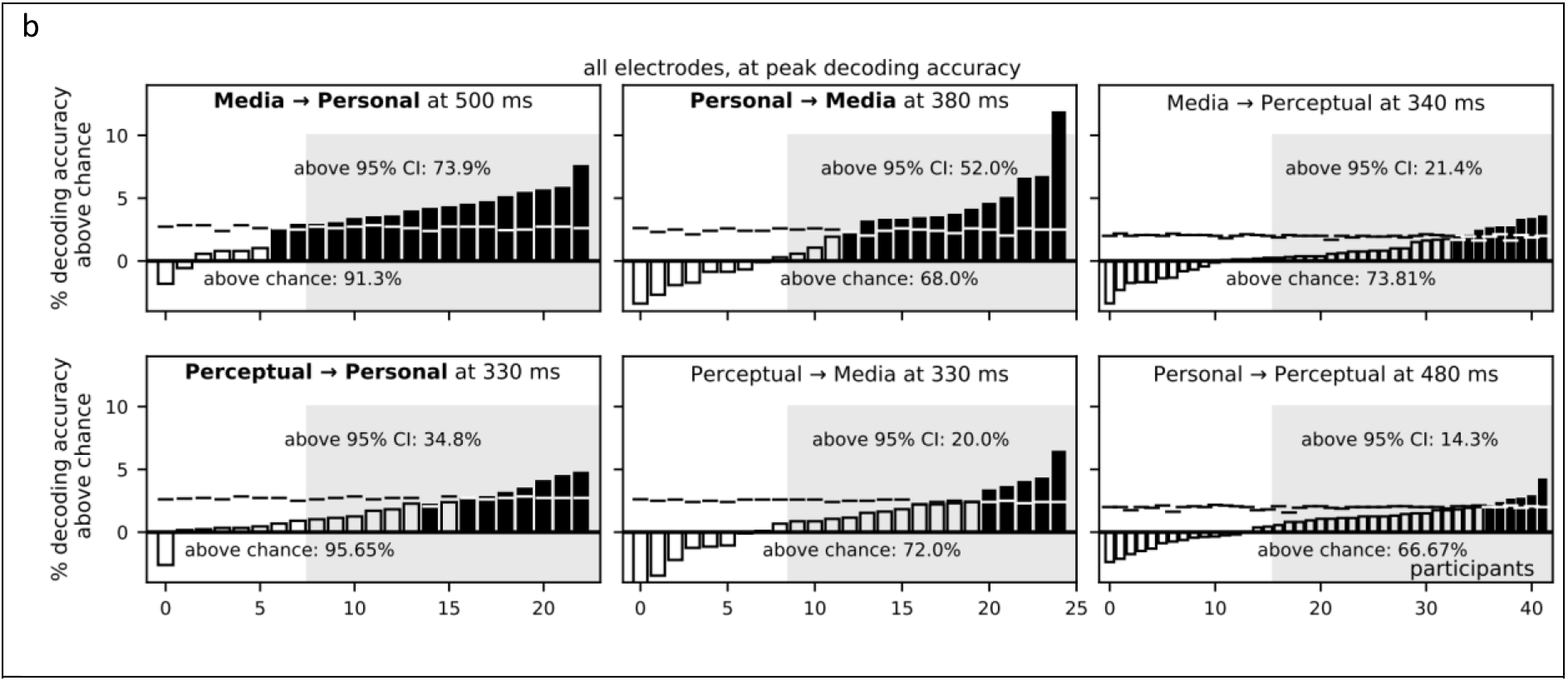
Cross-experiment familiarity decoding at a single participant-level, using bootstrapping analysis on decoding data from all electrodes. Each participant’s decoding accuracy is presented separately, with filled bars indicating values exceeding the bootstrapped 95% confidence interval (horizontal markers). (a) results of the overlapping 330 - 500 ms time windows (b) results at their respective peaks. Gray shaded areas denote the bootstrapped sample level 95% confidence intervals.

The temporal generalization analysis (**Figure 4., Supplementary Figure 2**.) revealed a sustained above-chance decoding accuracy starting around 170 ms, lasting consistently to around 600 ms or in some cases, beyond. Media predicted, between 180 ms to the end of the epoch from Personal between 80 ms onwards (cluster *p* = 0.0002). Reversely, Personal predicted, from 160 ms, Media, from 190 ms to 700 ms (cluster *p* = 0.017). Finally, Perceptual, between 170 and 1170 ms, predicted Personal, from 80 ms to 1010 ms, (cluster *p* = 0.019).

**Figure 4.**
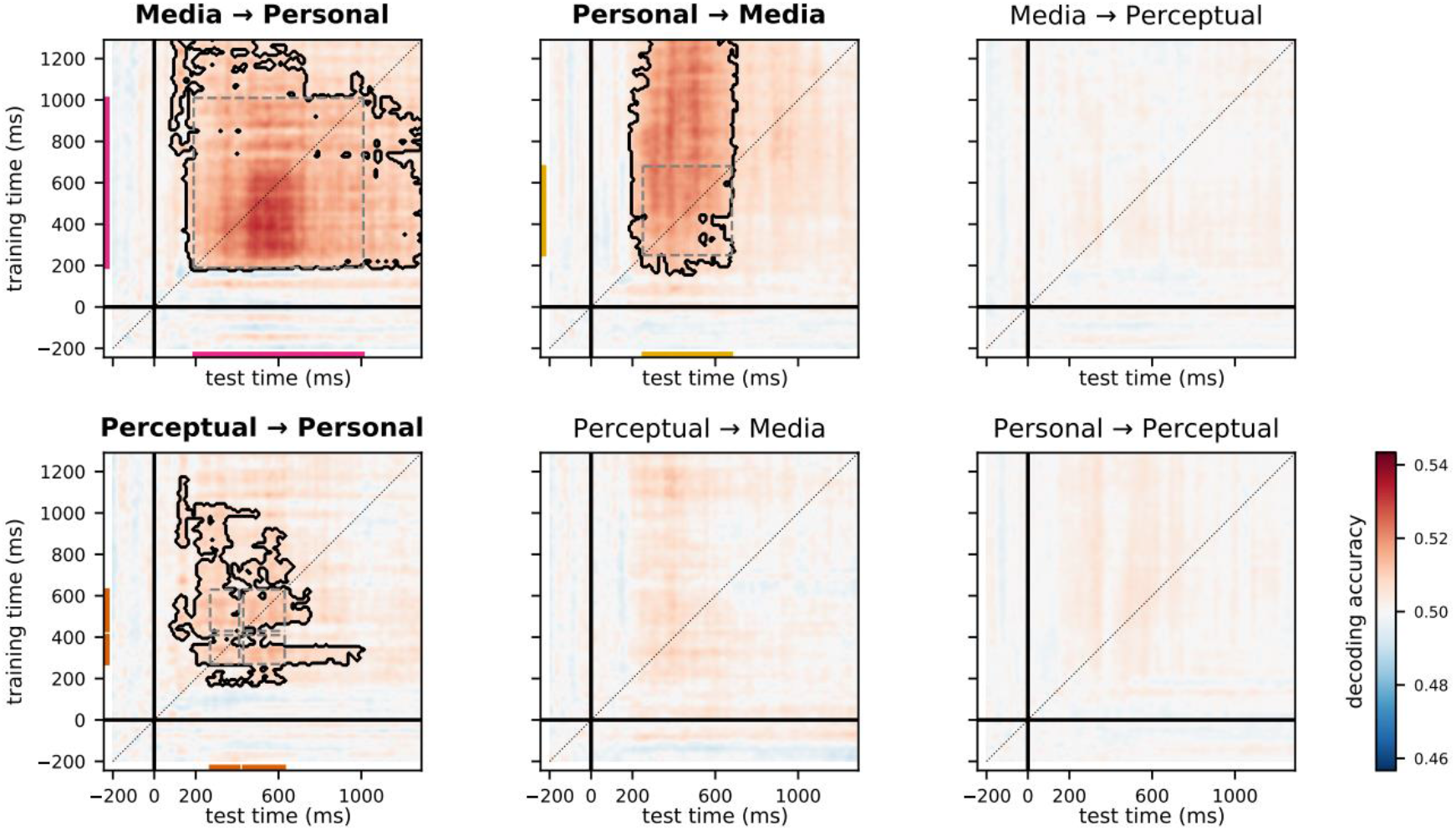
Cross-experiment temporal generalization analysis on all electrodes. For the results of the temporal generalization analyses separately in the different ROIs, see **Supplementary Figure 2**.

**Figure 5.**
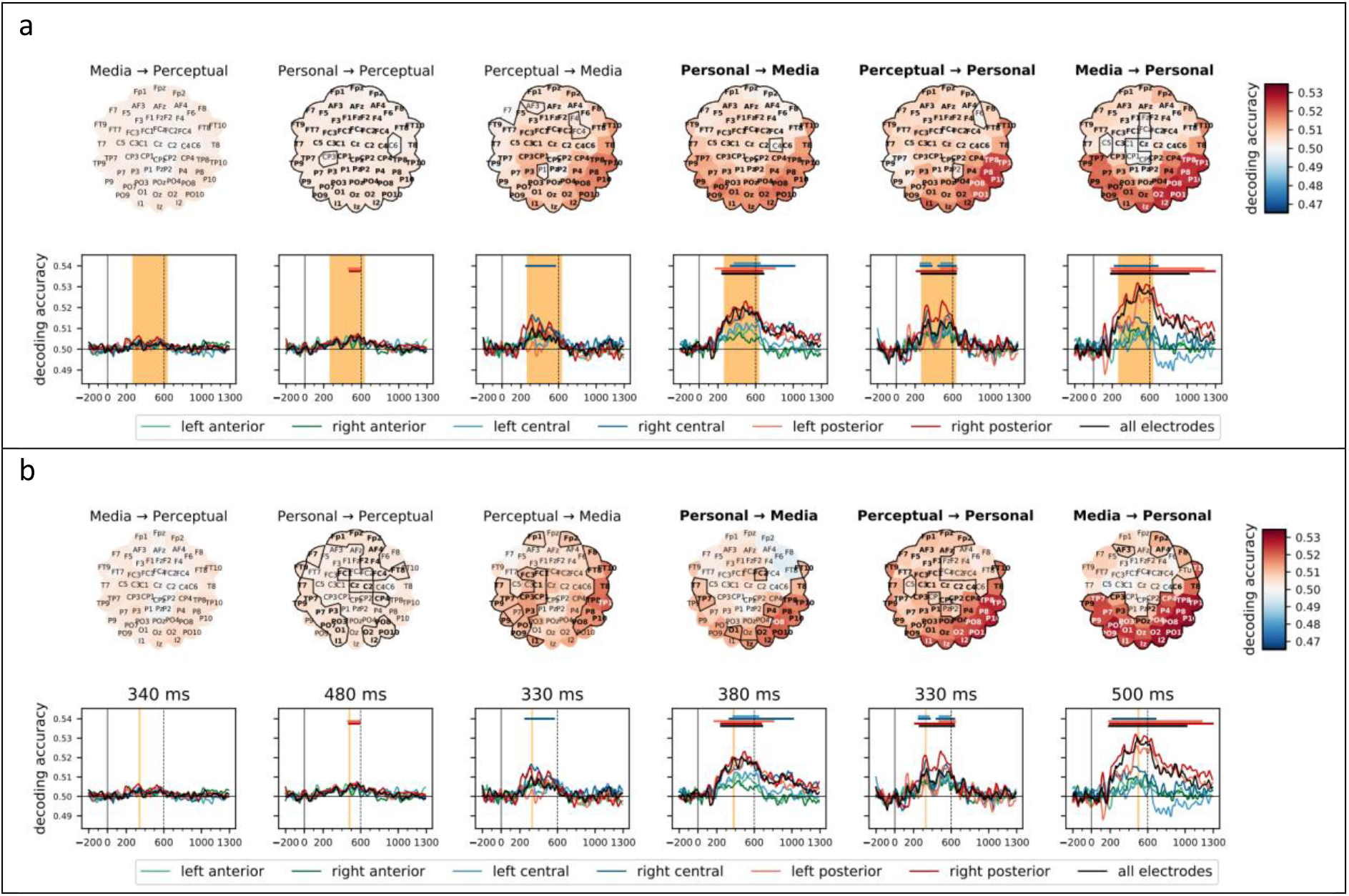
Region-of-interest and searchlight analyses. In three out of six train/test pairs involving all three experiments, significant decoding accuracies based on all electrodes (black horizontal marker) overlapped between 270 and 630 ms after stimulus onset (a). Further two train/test pairs revealed significant clusters at posterior or central ROIs within this interval. Peak decoding accuracies in all analyses (b) fell between this overlapping interval. The top panel scalp maps show the averaged searchlight-based decoding accuracies (electrodes belonging to significant spatio-temporal clusters marked) in the different train/test evaluations averaged over the overlapping timepoints. The bottom part shows the time-resolved ROI-based decoding results. Horizontal markers denote clusters with significantly different decoding accuracies against chance (two-tailed cluster permutation test, p < 0.05). For detailed results, see **Supplementary Figure 3**.

### Regions of Interest

To track representational organization across electrode space, we repeated the above analysis across the 6 electrode clusters also used in previous studies (Ambrus et al, 2019, 2021, in press). For further details see **Supplementary Figure 1** and **Supplementary Table 1**.

ROI analyses shown familiarity effects at posterior and central areas, mainly falling within the 300 to 600 ms interval. In Personal → Media, a right posterior cluster with an extent of 250 to 670 ms, peaking at 490 ms (peak *t*_24_ = 4.3, peak Cohen’s *d* = 0.86) two left posterior clusters at 180 – 340 ms with a peak at 270 ms (peak *t*_24_ = 2.83, peak Cohen’s *d* = 0.57) and 360 to 800 ms with a peak at 520 ms (peak *t*_24_ = 4.57, peak Cohen’s *d* = 0.91), a right central cluster between 340 and 1010 ms with a peak at 530 ms (peak *t*_24_ = 3.98, peak Cohen’s *d* = 0.80), and a left central cluster between 380 and 640 ms with a peak at 500 ms (peak *t*_24_ = 2.44, peak Cohen’s *d* = 0.59) emerged. In Media → Personal, a right posterior cluster between 200 and 1290 ms with a peak at 470 ms (peak *t*_22_ = 7.86, peak Cohen’s *d* = 1.64), three left posterior clusters, between 190 and 290 ms with a peak at 270 ms (peak *t*_22_ = 3.90, peak Cohen’s *d* = 0.82), between 320 and 990 ms with a peak at 480 ms (peak *t*_22_ = 4.96, peak Cohen’s *d* = 1.03), and a late 1010 to 1170 ms cluster with a peak at 1110 ms (peak *t*_22_ = 3.58, peak Cohen’s *d* = 0.75) and a right central cluster between 230 and 680 ms with a peak at 350 ms (peak *t*_22_ = 4.50, peak Cohen’s *d* = 1.04) were shown. In Perceptual → Personal, a right posterior cluster between 220 and 630 ms with a peak at 490 ms (peak *t*_22_ = 6.1, Cohen’s *d* = 1.27), a left posterior cluster between 480 and 640 ms with a peak at 520 ms (peak *t*_22_ = 5.29, Cohen’s *d* = 1.10), two right central clusters between 260 and 370 ms with a peak at 330 ms (peak *t*_22_ = 5.27, Cohen’s *d* = 1.10) and between 450 and 630 ms with a peak at 540 ms (peak *t*_22_ = 5.17, Cohen’s *d* = 1.08), two left central clusters between 260 and 360 ms with a peak at 330 ms (peak *t*_22_ = 3.58, Cohen’s *d* = 0.75) and 480 to 580 ms with a peak at 500 ms (peak *t*_22_ = 3.47, Cohen’s *d* = 0.72) emerged. In Personal → Perceptual, two posterior clusters, one on the right between 480 and 580 ms with a peak at 560 ms (peak *t*_41_ = 3.31, peak Cohen’s *d* = 0.51) and one on the left between 470 and 590 ms peaking at 520 ms (peak *t*_41_ = 3.12, peak Cohen’s *d* = 0.48) were found. In Perceptual → Media, a single right central cluster between 260 and 560 ms with a peak at 320 ms (peak *t*_24_ = 3.59, peak Cohen’s *d* = 0.72) was observed. In Media → Perceptual no significant effect was found.

### Searchlight

Apart from Media → Perceptual, extensive significant spatio-temporal clusters, encoding familiarity, were observed in several searchlight analyses. In Media → Personal a significant cluster (cluster *p* < 0.001), starting from 180 ms at bilateral temporo-parieto-occipital electrodes, was observed. Also starting at 180 ms (electrode I1), there was a significant cluster found in Personal → Media (cluster *p* = 0.014), lasting until 700 ms after stimulus onset. A significant cluster with an even earlier onset (100 ms, at Oz and POz) was observed in Perceptual → Personal (cluster *p* < 0.001), lasting until 1210 ms. Between 210 and 680 ms, Perceptual → Media (cluster *p* = 0.01, starting at PO8 and P4) and between 410 and 870 ms, Personal → Perceptual (starting at FT9, cluster *p* = 0.007) also yielded significant clusters.

Analysis of the searchlight results revealed peak decoding accuracies at right parieto-occipito-temporal sites in all evaluations. In both Media → Personal and Personal → Media, maximum decoding accuracy was found at PO8, peaking at 560 ms (peak Cohen’s *d* = 1.83) and 490 ms (peak Cohen’s *d* = 0.89), respectively. In Perceptual → Personal the peak decoding accuracy was at P8, with a peak at 480 ms (peak Cohen’s *d* = 1.55). In Perceptual → Media, peak decoding accuracy was observed at TP10 at 370 ms (peak Cohen’s *d* = 0.85). In Personal → Perceptual, peak decoding accuracy was found at P8 at 560 ms (peak Cohen’s *d* = 0.61), and in Media → Perceptual at I2 at 330 ms (peak Cohen’s *d* = 0.32). **Figure 6**. shows the scalp distributions in the six cross-experiment searchlight decoding evaluations at and the bootstrap-based subject-level reliability calculations at their respective peak decoding latencies. For further details see **Supplementary Figure 3**.

**Figure 6.**
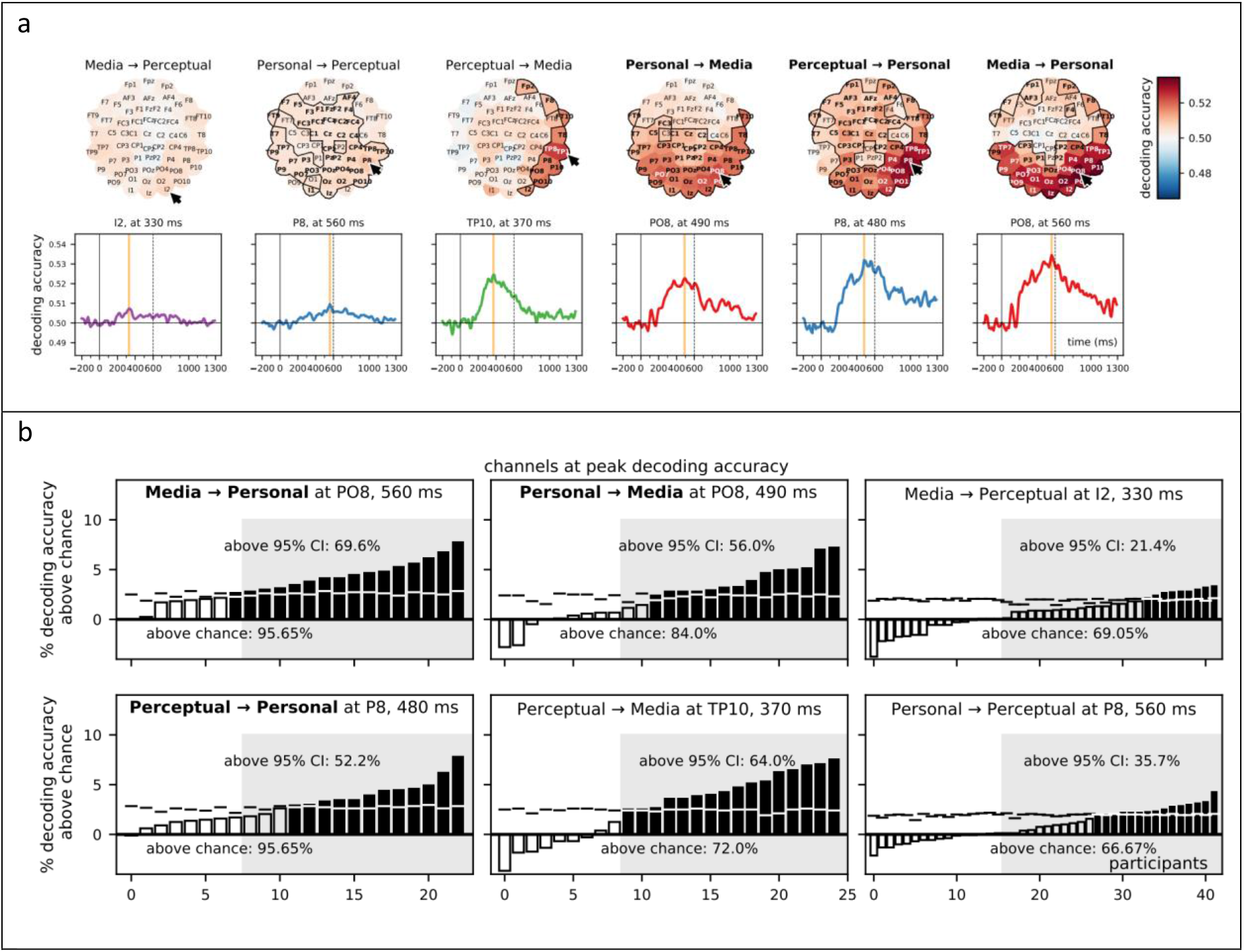
Cross-experiment searchlight analysis. Electrodes with the highest peak decoding accuracy at their respective peak latencies. (a) Highest peak decoding accuracies are seen over the right-hemisphere occipito-temporo-parietal electrodes (black arrows), ranging from 330 to 560 ms. (b) Bootstrapping analyses on the data of these electrodes at peak accuracy times revealed above-chance decoding in the majority of the participants in every cross-experiment combination. The decoding accuracy of individual participants is shown, with filled bars indicating values exceeding the bootstrapped 95% confidence interval (horizontal markers). Gray shaded areas denote the bootstrapped sample level 95% confidence intervals. Pairs with significant clusters in decoding from all electrodes are highlighted in bold typeface.

## Discussion

The current analysis is based on data from Ambrus et al., 2021 (in press) which explored the temporal dynamics of emerging face identity and familiarity information under different of familiarization conditions. Time-resolved representational similarity analysis of EEG data revealed that the level of familiarization has a robust effect on familiarity representations: they are strongly visible over posterior and central regions following personal, somewhat weaker after media familiarization, and they are absent following perceptual familiarization.

In this present study we set out to uncover the existence of shared face familiarity signal. We applied a cross-experiment decoding approach where evoked responses from participants in one experiment are decoded using classifiers trained on the aggregated data of the participants of another study. It is important to note that three experiments of the current study differed in (1) the mode and length of familiarization, (2) the stimulus material, (3) the number of images of a given identity, (4) the task, and (5) the participants tested.

Despite all these substantial differences between the three experiments, we found significant sustained cross-experiment familiarity decoding accuracies, most prominently in the Media ↔ Personal, and the Perceptual → Personal familiarization conditions, overlapping between 270-630 ms. Bootstrapping analyses within the peak accuracy intervals (330 – 500 ms) have shown above-chance decoding in most of the participants, albeit the participant-level reliability did not reach the sample-level 95% confidence interval (Figure 3a). Additional ROI-and searchlight-based analyses revealed bilateral posterior and central effects, with a more pronounced right-hemisphere weight.

We observed the most evident cross-experiment familiarity effect between the Media and Personal familiarization experiments, where the depth of familiarization was arguably the highest. In the Media experiment, in the span of two weeks, participants viewed a season of a television series, while in the Personal experiment, participants interacted with the experimentally familiarized identities personally for ca. 1 hour on three consecutive days. Somewhat unexpectedly, a similar effect emerged in Perceptual → Personal albeit more restricted in time. This is surprising as no significant familiarity effect was observed in the Perceptual experiment, in which identities were familiarized through a very short sequential sorting task (based on Andrews et al. (2015), as implemented in Ambrus et al. (2017b)). Although no further significant decoding effects on all electrodes were found, in Perceptual → Media a right-central cluster with an onset and duration similar to other train/test pairs was observed, while in Personal → Perceptual a small bilateral posterior effect was seen, also falling within the overlapping time windows of the strongest effects found in other decoding pairs. Finally, no familiarity-related cross-experiment decoding effects were observed in Media → Perceptual.

There was some variation in the onset and duration of the effects, with searchlight and temporal generalization analyses flagging time points as early as ca. 100 and 160 ms, and ROI analyses showing onsets at around 180 – 200 ms in some cases. Literature regarding early effects is somewhat inconsistent (Ramon and Gobbini, 2018) – while some reported no familiarity-related modulation in such early phases (Huang et al., 2017), ERP amplitude increase (e.g. (Kloth et al., 2006)), *and* decrease (Todd et al., 2008) have also been described.

Recently, In an MEG-MVPA experiment Dobs et al. (2019) found a familiarity-related component about 400 to 600 ms after stimulus onset, using familiar and unfamiliar celebrities as stimuli. The authors hypothesized that this identity-independent familiarity effect may reflect memory-related activations or affective responses. Wiese et al. (2019) also described an ERP component elicited by highly familiar versus unfamiliar faces. This sustained familiarity effect (SFE) started at around 200 ms after stimulus onset and reached its maximum between 400 and 600 ms as well. The source of this effect is theorized to lie within the ventral visual pathway, and its magnitude was found to be modulated by the level of familiarity (larger for contrasts with personally familiar faces compared to lecturers or celebrities). This led the authors to suggest that the SFE may be driven by the integration of perceptual and affective information. Finally, Karimi-Rouzbahan et al. (2020) also observed a familiarity-related effect starting around 200 ms and peaking after ca. 400, with the highest ERP amplitude elicited by the participants own face, followed by personally familiar, celebrity, and unfamiliar faces. The most stable cross-experiment effects were also observed in the 400 – 600 ms time-window, with peak decoding accuracies between 330 - 500 ms, supporting these above-mentioned results. Based on their temporal and spatial characteristics, we suggest that the familiarity effects, reported by the previous studies and the cross-experiment decoding of the current study are related, and are indicative of a robust and general familiarity effect, which is largely independent of participants, experimental stimuli, and the method of familiarization.

Where significant familiarity-related effects were present, temporal generalization analyses on all electrodes revealed a sustained above-chance decoding performance starting around 200 ms, lasting consistently to around 600 ms or in some cases, beyond. This suggests that the classifiers picked up on features generated by a single process, maintained over time (King and Dehaene, 2014).

At this point, we can only speculate as to the nature of processing underlying this observed effect. Although it is difficult to make inferences regarding the anatomical structures generating these signals, its occipito-temporal dominance is consistent with sources reported in relation to the ventral visual pathway, and parts of the core and extended face networks are most certainly involved (Gobbini and Haxby, 2007; Kovács, 2020). The occipital face area (OFA), an early face-selective region that has been shown to take part in image-dependent lower-level feature analysis (Pitcher et al., 2007) is already differentially responsive to familiar and unfamiliar identities (Jonas et al., 2014; Ambrus et al., 2017a, 2019a; Amado et al., 2018), and may also play a causal role in the learning of new faces (Ambrus et al., 2017b). The fusiform face area (FFA) shows image-independent processing of face-information, it is therefore highly probable that its normal functioning is central to the recognition of familiar identities (Kanwisher et al., 1997). Finally, the superior temporal sulcus (STS) is more sensitive to dynamic/changeable visual features such as facial expressions (Zhang et al., 2016; Direito et al., 2019) (but see also (Calder and Young, 2016)) that may also carry diagnostic, identity-specific information, thus boosting familiarity signals. (Also worth noting here that face-responsive cells in the monkey superior temporal polysensory area codes for familiarity and social status (Young and Yamane, 1992)).

The relatively late peak decoding latency we observed may be an indication that it depends, at least partially, on post-perceptual processing that is fed back from more modality-independent, memory-related cortical areas. For example, regions considered to be outside the core face networks, such as different extrastriate visual areas, the structures in the anterior and medial temporal lobe (MTL) have been reliably demonstrated to exhibit stronger responses to familiar stimuli, including faces. The anterior temporal lobe (ATL), considered to be a semantic hub (Anzellotti, 2017), is involved in associations of person-related semantics (Morton et al., 2021), e.g. a connection between a name and a face (Tsukiura et al., 2010). Structures within the MTL (e.g. the amygdala, perirhinal cortex and hippocampus) are involved in the storing and processing of biographical/autobiographical information related to faces, including the feelings of familiarity, and respond abruptly when sufficient information for familiar face recognition is accumulated (Ramon et al., 2015).

The fact that Personal (and to some extent, Media) was decodable from Perceptual suggests that seeds of this effect are already present in purely image-based individuation suggests that seeds of this effect are already present in purely image-based individuation. s This might be an indication that the information is “present” enough to be picked up by classifiers at training, but drowned out by irrelevant features or “noise” in decoding, as generalization from the dataset with a small number of informative features to the dataset with a large number of informative features gives better classification performance than vice versa (van den Hurk and Op de Beeck, 2019). This seems to imply that contributions from semantic, autobiographical, and affective processes are less essential, as these facets played little role in the Perceptual familiarization experiment, thus memory processes related to face-feature-based recall might be more promising candidates. Still, it is possible that even for short term, perceptual familiarization the whole cascade is engaged, for example because even in this case the familiarity signal triggers search processes in long term storage and/or semantic and affective hubs. Further research is needed to precisely pin down the exact cognitive machinery behind our results. For example, methods such as M/EEG-fMRI fusion (Cichy and Oliva, 2020; Muukkonen et al., 2020), combining the spatial and temporal resolution of these techniques, may give us further insight.

Finally, we need to stress that our findings by no means imply that familiarity information first becomes available at this late time interval. As we have seen, familiarity information was present in time points as early as 100 ms post stimulus presentation in some clusters. Furthermore, Dobs et al. demonstrated differential face-sex read-out in a cluster containing time points even as early as 100 ms post stimulus onset. This indicates that familiarity with a face may influence processing much earlier, thus the information can be available to at least some of the systems involved. It has been demonstrated that unique configuration of familiar facial features may influence early perceptual processing even at the V1 level. It is thus conceivable that higher areas of the core and extended face networks feed the outcome of their analyses back to early visual areas, allowing for a facilitated detection without an explicit recognition, a quasi “pre-recognition” (Visconti di Oleggio Castello et al., 2014) of highly familiar faces. Although independent confirmation of this hypothesis is required, the cross-temporal decodability of the early and later signals suggests that at least some of the coding may be shared even in the earliest stages of face processing.

On a methodological note, this study has shown that cross experiment decoding is uniquely suited for finding general neural patterns that are indicative of a shared processing of information. It has the potential to be more sensitive than within-subject decoding, as it benefits from more training data and is less confounded by idiodincratic participant-level effects and stimulus properties. We hope to see this method being utilized to advance our understanding of brain dynamics in the future.

## Summary

In conclusion, we confirmed the existence of a neural signature of face familiarity using cross-experiment decoding of event-related potentials evoked by unknown and experimentally familiarized faces from a set of experiments involving different participants, stimuli, and modes of familiarization. This effect is remarkably similar in temporal and spatial characteristics to electrophysiological components reported recently, suggesting that the same effect was observed in these previous studies. The fact that familiarity was found to be decodable between participants and experiments makes this component a strong candidate for a general, automatic neural signature of face familiarity. Furthermore, the sustained pattern of temporal generalization suggests a single underlying information processing step that is maintained across time. Future studies need to elucidate the precise functional-neuroanatomical underpinnings of this process.

## Supplementary Material

**Supplementary Figure 1.** Plots showing the results of the time-resolved cross-experiment decoding analysis on pre-defined regions of interests.

**Supplementary Table 1.** Statistics on the results of the time-resolved cross-experiment decoding analysis on pre-defined regions of interests.

**Supplementary Figure 2.** Results of the cross-experiment temporal generalization analyses

**Supplementary Figure 3.** Results of the searchlight cross-experiment decoding analyses

## References

Ambrus GG, Eick CM, Kaiser D, Kovács G (2021, in press) Getting to know you: emerging neural representations during face familiarization. Accepted in The Journal of Neuroscience.

Amado C, Kovács P, Mayer R, Ambrus GG, Trapp S, Kovács G (2018) Neuroimaging results suggest the role of prediction in cross-domain priming. Sci Rep 8:10356 Available at: http://www.nature.com/articles/s41598-018-28696-0 [Accessed July 9, 2018].

Ambrus GG, Amado C, Krohn L, Kovács G (2019a) TMS of the occipital face area modulates cross-domain identity priming. Brain Struct Funct 224:149–157 Available at: http://link.springer.com/10.1007/s00429-018-1768-0.

Ambrus GG, Dotzer M, Schweinberger SR, Kovács G (2017a) The occipital face area is causally involved in the formation of identityspecific face representations. Brain Struct Funct.

Ambrus GG, Kaiser D, Cichy RM, Kovács G (2019b) The Neural Dynamics of Familiar Face Recognition. Cereb Cortex 29:4775–4784 Available at: https://dx.doi.org/10.1093/cercor/bhz010.

Ambrus GG, Windel F, Burton AM, Kovács G (2017b) Causal evidence of the involvement of the right occipital face area in face-identity acquisition. Neuroimage 148:212–218 Available at: https://www.sciencedirect.com/science/article/pii/S1053811917300514.

Andrews S, Jenkins R, Cursiter H, Burton AM, Andrews S, Jenkins R, Cursiter H, Telling AMB, Andrews S, Jenkins R, Cursiter H, Burton AM (2015) Telling faces together: Learning new faces through exposure to multiple instances Telling faces together: Learning new faces through exposure to multiple instances. Q J Exp Psychol 0218:2041–2050 Available at: https://www.ncbi.nlm.nih.gov/pubmed/25607814.

Anzellotti S (2017) Anterior temporal lobe and the representation of knowledge about people. Proc Natl Acad Sci U S A 114:4042–4044.

Barragan-Jason G, Cauchoix M, Barbeau EJ (2015) The neural speed of familiar face recognition. Neuropsychologia 75:390–401.

Buttle H, Raymond JE (2003) High familiarity enhances visual change detection for face stimuli. Percept Psychophys 65:1296–1306.

Caharel S, Poiroux S, Bernard C, Thibaut F, Lalonde R, Rebai M (2002) ERPs associated with familiarity and degree of familiarity during face recognition. Int J Neurosci 112:1499–1512.

Calder AJ, Young AW (2016) Understanding the recognition of facial identity and facial expression. Facial Expr Recognit Sel Work Andy Young 6:41–64.

Cichy RM, Oliva A (2020) A M/EEG-fMRI Fusion Primer: Resolving Human Brain Responses in Space and Time. Neuron 107:772–781.

Debruille JB, Guillem F, Renault B (1998) ERPs and chronometry of face recognition: Following-up Seeck et al. and George et al. Neuroreport 9:3349–3353 Available at: http://www.ncbi.nlm.nih.gov/pubmed/9855278 [Accessed July 7, 2018].

Di Nocera F, Ferlazzo F (2000) Resampling approach to statistical inference: Bootstrapping from event-related potentials data. Behav Res Methods, Instruments, Comput 32:111–119.

Direito B, Lima J, Simões M, Sayal A, Sousa T, Lührs M, Ferreira C, Castelo-Branco M (2019) Targeting dynamic facial processing mechanisms in superior temporal sulcus using a novel fMRI neurofeedback target. Neuroscience 406:97–108.

Dobs K, Isik L, Pantazis D, Kanwisher N (2019) How face perception unfolds over time. Nat Commun 10.

Duchaine B, Yovel G (2015) A Revised Neural Framework for Face Processing. Annu Rev Vis Sci 1:393–416.

Ellis HD, Lewis MB (2001) Capgras delusion: A window on face recognition. Trends Cogn Sci 5:149–156.

Gobbini MI, Haxby J V. (2007) Neural systems for recognition of familiar faces. Neuropsychologia 45:32–41 Available at: https://www.ncbi.nlm.nih.gov/pubmed/16797608.

Gramfort A, Luessi M, Larson E, Engemann DA, Strohmeier D, Brodbeck C, Goj R, Jas M, Brooks T, Parkkonen L, Hämäläinen M (2013) MEG and EEG data analysis with MNE-Python. Front Neurosci Available at: https://www.frontiersin.org/articles/10.3389/fnins.2013.00267/full.

Grootswagers T, Wardle SG, Carlson TA (2017) Decoding dynamic brain patterns from evoked responses: A tutorial on multivariate pattern analysis applied to time series neuroimaging data. J Cogn Neurosci 29:677–697 Available at: https://www.mitpressjournals.org/doi/abs/10.1162/jocn_a_01068.

Huang W, Wu X, Hu L, Wang L, Ding Y, Qu Z (2017) Revisiting the earliest electrophysiological correlate of familiar face recognition. Int J Psychophysiol 120:42–53.

Johnston P, Overell A, Kaufman J, Robinson J, Young AW (2016) Expectations about person identity modulate the face-sensitive N170. Cortex 85:54–64.

Johnston RA, Edmonds AJ (2009) Familiar and unfamiliar face recognition: A review. Memory 17:577–596 Available at: http://www.ncbi.nlm.nih.gov/pubmed/19548173.

Jonas J, Rossion B, Krieg J, Koessler L, Colnat-Coulbois S, Vespignani H, Jacques C, Vignal JP, Brissart H, Maillard L (2014) Intracerebral electrical stimulation of a face-selective area in the right inferior occipital cortex impairs individual face discrimination. Neuroimage 99:487–497.

Kaiser D, Oosterhof NN, Peelen M V. (2016) The Neural Dynamics of Attentional Selection in Natural Scenes. J Neurosci 36:10522–10528 Available at: http://www.jneurosci.org/cgi/doi/10.1523/JNEUROSCI.1385-16.2016.

Kanwisher NG, McDermott J, Chun MM (1997) The fusiform face area: a module in human extrastriate cortex specialized for face perception. J Neurosci 17:4302–4311.

Kaplan JT, Man K, Greening SG (2015) Multivariate cross-classification: Applying machine learning techniques to characterize abstraction in neural representations. Front Hum Neurosci 9.

Karimi-Rouzbahani H, Remezani F, Woolgar A, Rich A, Ghodrati M (2020) Perceptual difficulty modulates the direction of information flow in familiar face recognition. bioRxiv:2020.08.10.245241 Available at: https://doi.org/10.1101/2020.08.10.245241.

King JR, Dehaene S (2014) Characterizing the dynamics of mental representations: The temporal generalization method. Trends Cogn Sci 18:203–210.

Kloth N, Dobel C, Schweinberger SR, Zwitserlood P, Bölte J, Junghöfer M (2006) Effects of personal familiarity on early neuromagnetic correlates of face perception. Eur J Neurosci 24:3317–3321.

Kovács G (2020) Getting to know someone: Familiarity, person recognition, and identification in the human brain. J Cogn Neurosci 32:2205–2225.

Landi SM, Freiwald WA (2017) Two areas for familiar face recognition in the primate brain. Science (80-) 357:591–595.

Millen AE, Hancock PJB (2019) Eye see through you! Eye tracking unmasks concealed face recognition despite countermeasures. Cogn Res Princ Implic 4.

Morton NW, Zippi EL, Noh S, Preston AR (2021) Semantic knowledge of famous people and places is represented in hippocampus and distinct cortical networks. J Neurosci.

Muukkonen I, Ölander K, Numminen J, Salmela VR (2020) Spatio-temporal dynamics of face perception. Neuroimage 209.

Pedregosa F, Varoquaux G, Gramfort A, Michel V, Thirion B, Grisel O, Blondel M, Prettenhofer P, Weiss R, Dubourg V, others (2011) Scikit-learn: Machine learning in Python. J Mach Learn Res 12:2825–2830.

Persike M, Meinhardt-Injac B, Meinhardt G (2013) The preview benefit for familiar and unfamiliar faces. Vision Res 87:1–9.

Pitcher D, Walsh V, Yovel G, Duchaine B (2007) TMS evidence for the involvement of the right occipital face area in early face processing. Curr Biol 17:1568–1573.

Ramon M, Gobbini MI (2018) Familiarity matters: A review on prioritized processing of personally familiar faces. Vis cogn 26:179–195.

Ramon M, Vizioli L, Liu-Shuang J, Rossion B (2015) Neural microgenesis of personally familiar face recognition. Proc Natl Acad Sci 112:E4835–E4844 Available at: http://www.pnas.org/lookup/doi/10.1073/pnas.1414929112.

Rosenzweig G, Bonneh YS (2019) Familiarity revealed by involuntary eye movements on the fringe of awareness. Sci Rep 9.

Sanchez JFQ, Liu X, Zhou C, Hildebrandt A (2021) Nature and Nurture Shape Structural Connectivity in the Face Processing Brain Network. Neuroimage:117736.

Todd RM, Lewis MD, Meusel LA, Zelazo PD (2008) The time course of social-emotional processing in early childhood: ERP responses to facial affect and personal familiarity in a Go-Nogo task. Neuropsychologia 46:595–613.

Tranel D, Damasio AR (1985) Knowledge without awareness: An autonomic index of facial recognition by prosopagnosics. Science (80-) 228:1453–1454.

Tsukiura T, Mano Y, Sekiguchi A, Yomogida Y, Hoshi K, Kambara T, Takeuchi H, Sugiura M, Kawashima R (2010) Dissociable roles of the anterior temporal regions in successful encoding of memory for person identity information. J Cogn Neurosci 22:2226–2237.

van den Hurk J, Op de Beeck HP (2019) Generalization asymmetry in multivariate cross-classification: When representation A generalizes better to representation B than B to A. bioRxiv.

Verhallen RJ, Bosten JM, Goodbourn PT, Lawrance-Owen AJ, Bargary G, Mollon JD (2016) General and specific factors in the processing of faces. Vision Res.

Virtanen P et al. (2020) SciPy 1.0: fundamental algorithms for scientific computing in Python. Nat Methods 17:261–272.

Visconti di Oleggio Castello M, Guntupalli JS, Yang H, Gobbini MI (2014) Facilitated detection of social cues conveyed by familiar faces. Front Hum Neurosci 8.

Wiese H, Tüttenberg SC, Ingram BT, Chan CYX, Gurbuz Z, Burton AM, Young AW (2019) A Robust Neural Index of High Face Familiarity. Psychol Sci 30:261–272.

Wilmer JB, Germine L, Chabris CF, Chatterjee G, Williams M, Loken E, Nakayama K, Duchaine B (2010) Human face recognition ability is specific and highly heritable. Proc Natl Acad Sci U S A 107:5238–5241 Available at: http://www.pnas.org/cgi/doi/10.1073/pnas.0913053107.

Yan X, Young AW, Andrews TJ (2017) The automaticity of face perception is influenced by familiarity. Attention, Perception, Psychophys 79:2202–2211.

Young MP, Yamane S (1992) Sparse population coding of faces in the intertemporal cortex. Science (80-) 256:1327–1331.

Zhang H, Japee S, Nolan R, Chu C, Liu N, Ungerleider LG (2016) Face-selective regions differ in their ability to classify facial expressions. Neuroimage 130:77–90.

