## Supplementary figures and images for "Evidence for a general neural signature of face familiarity"

### Supplementary Figure 1

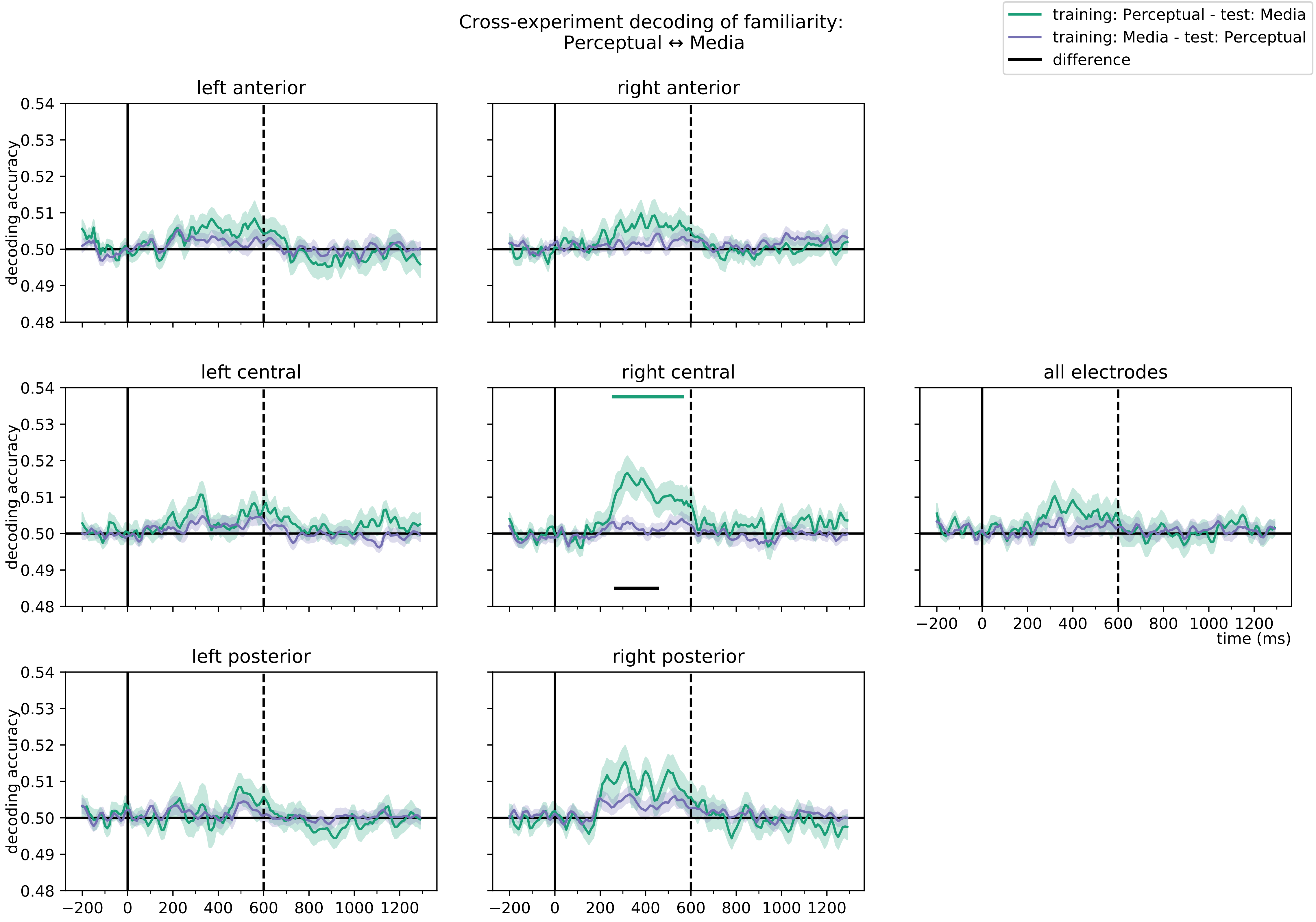

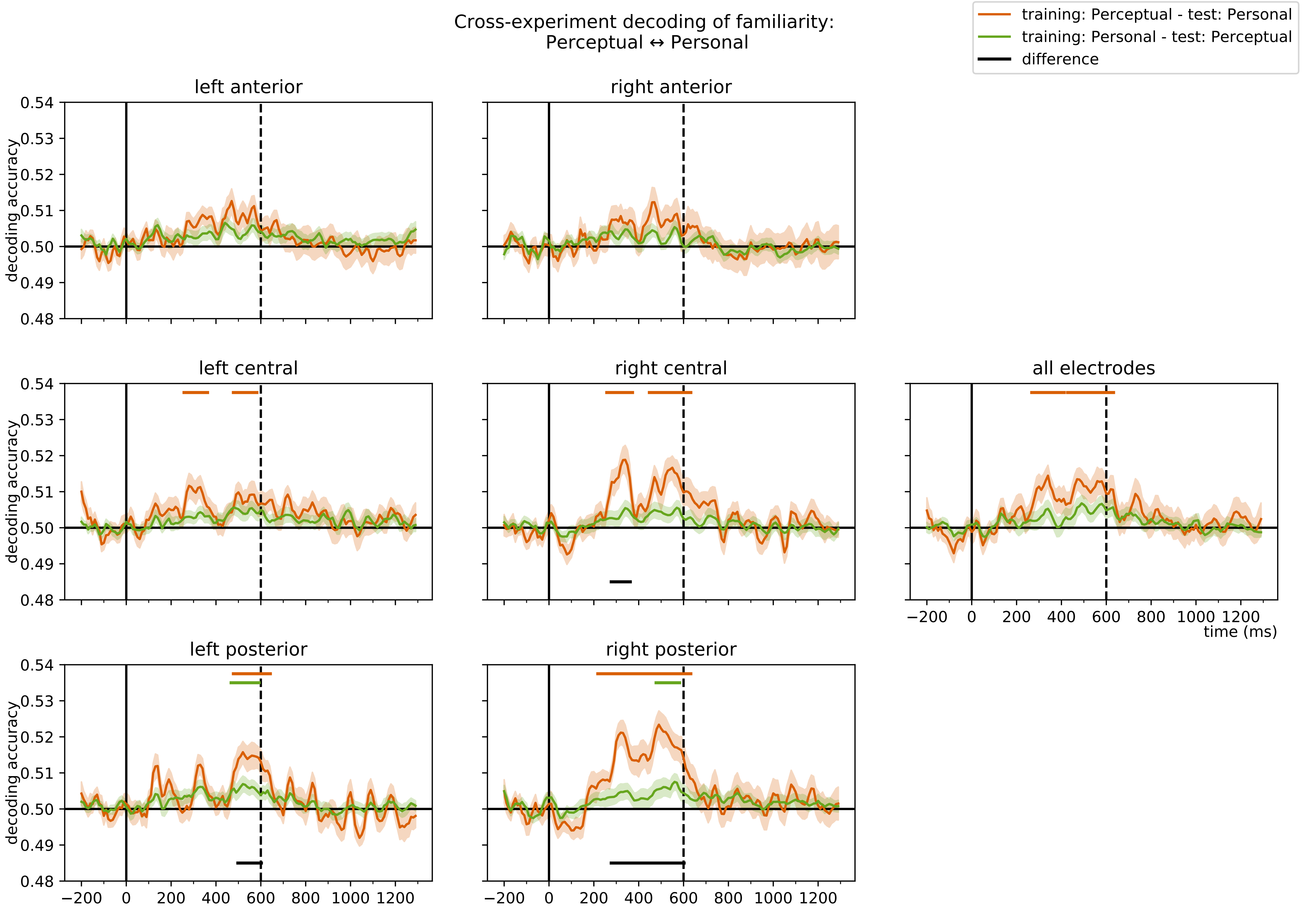

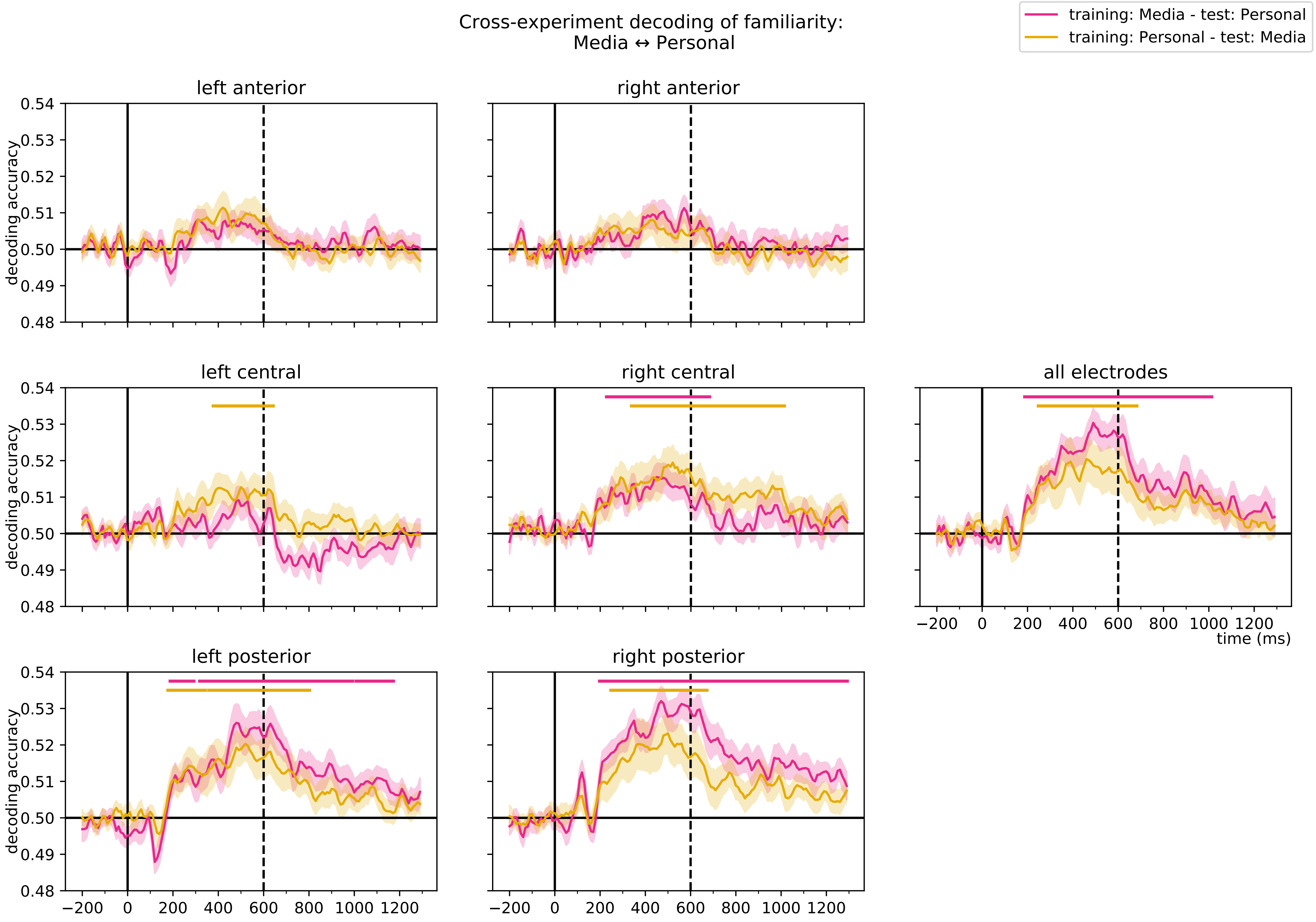

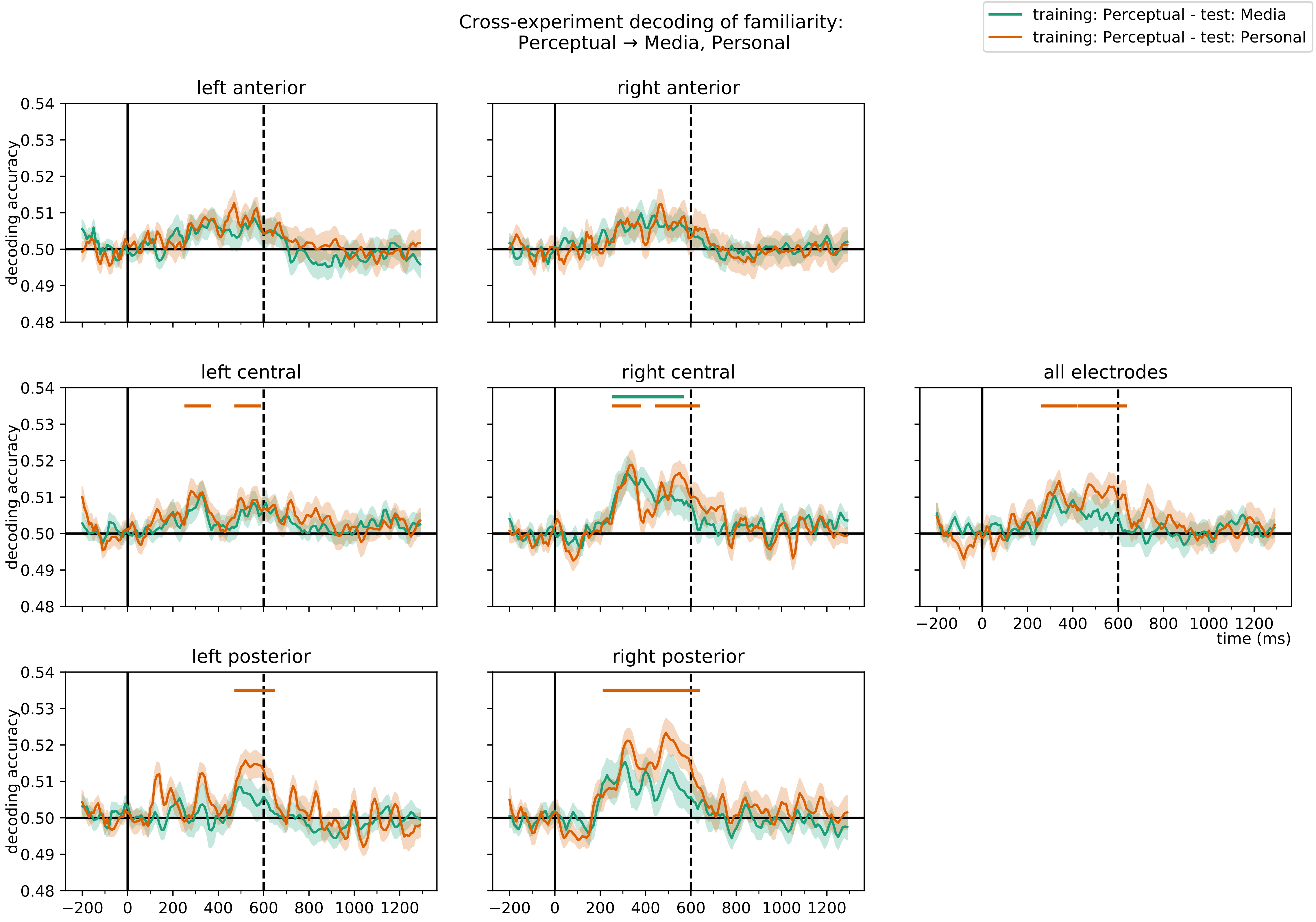

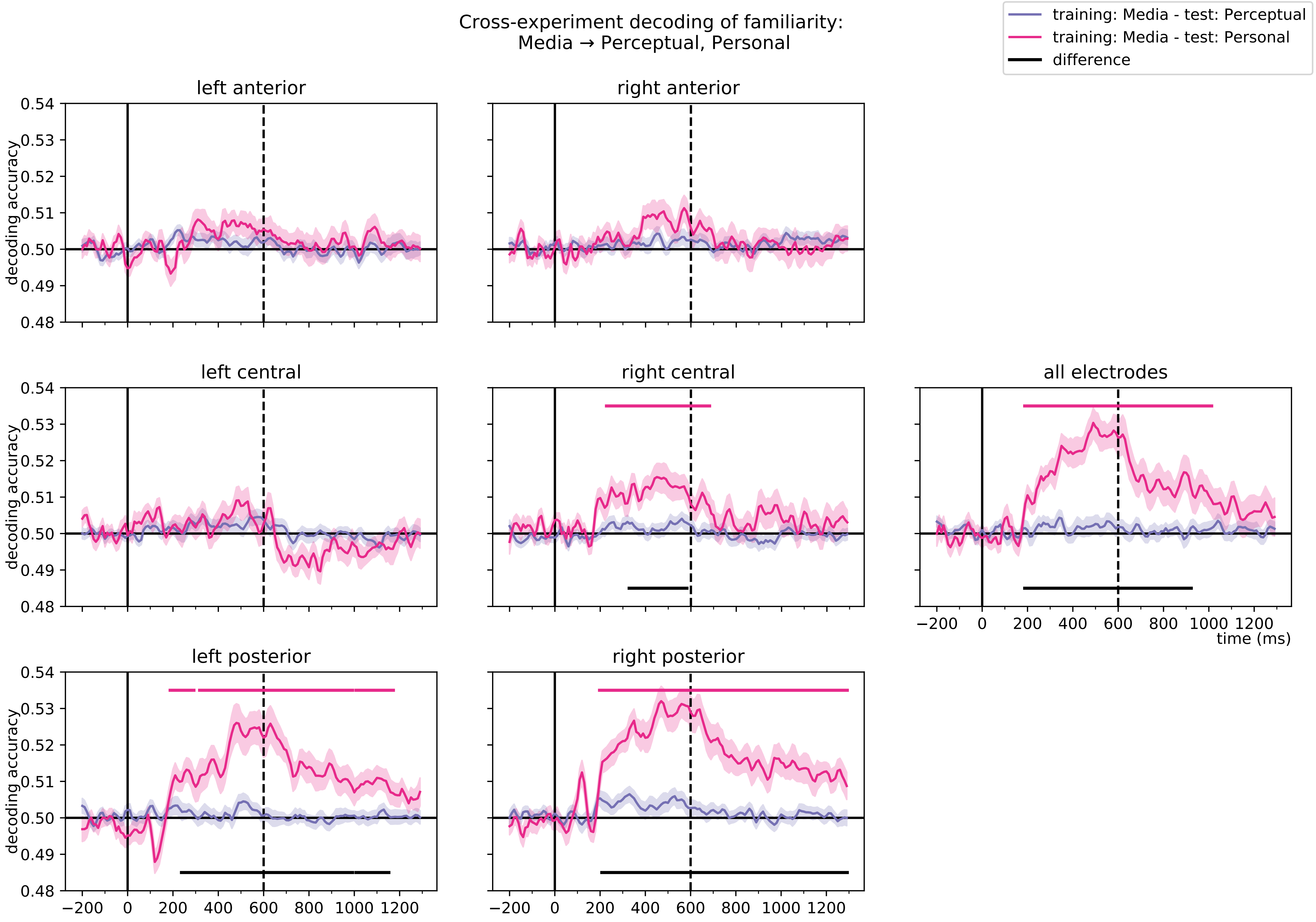

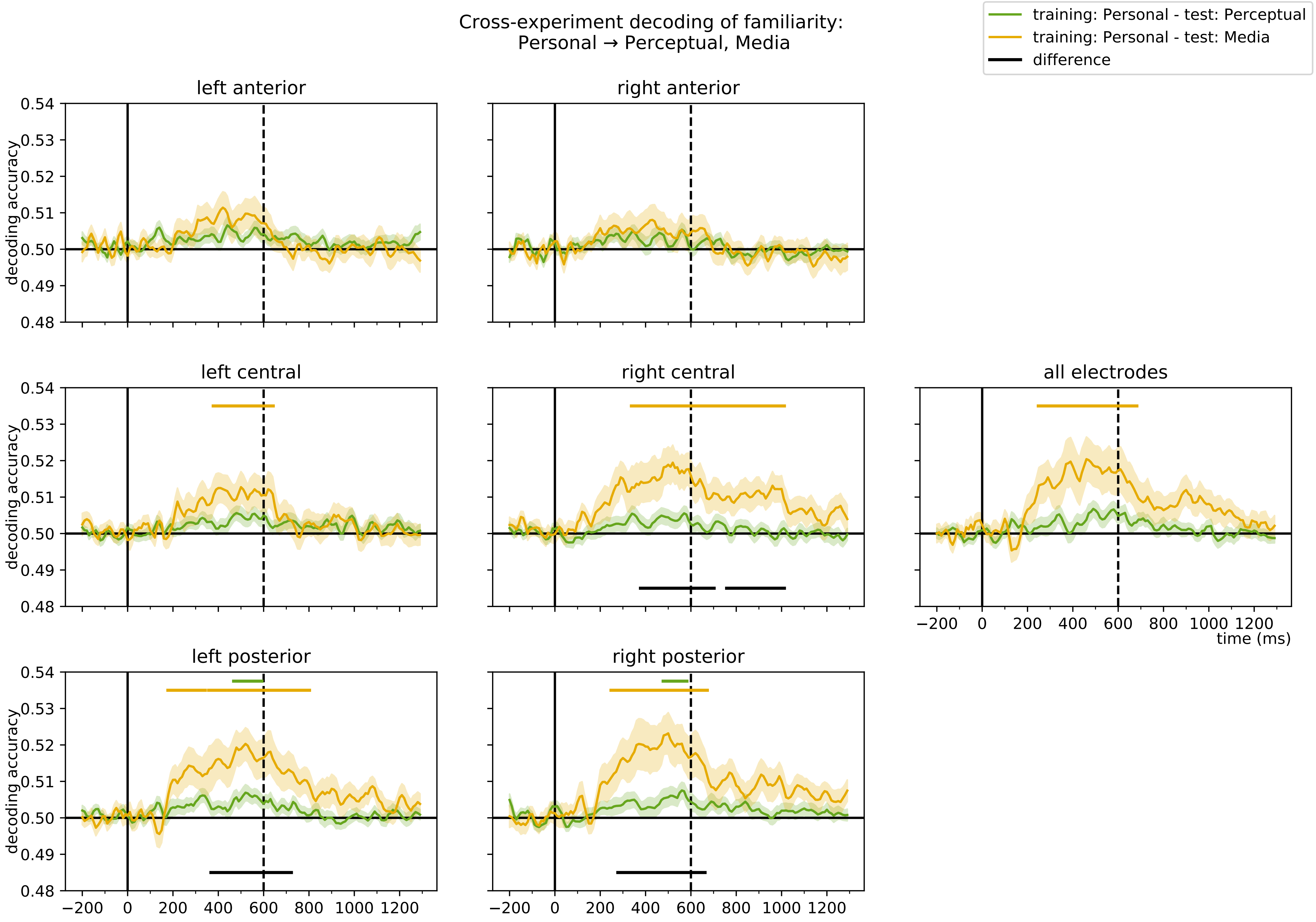

### Supplementary Figure 2

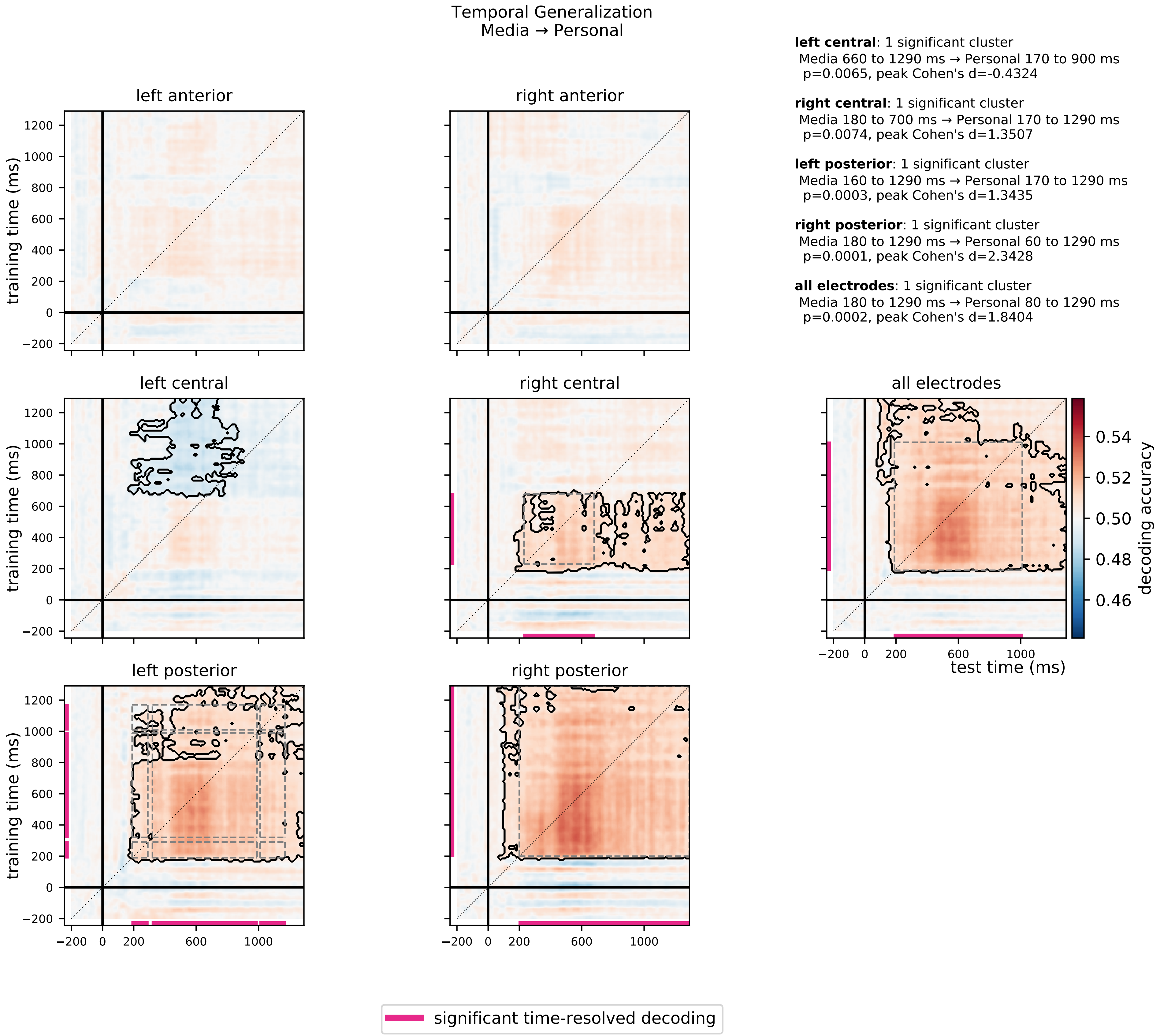

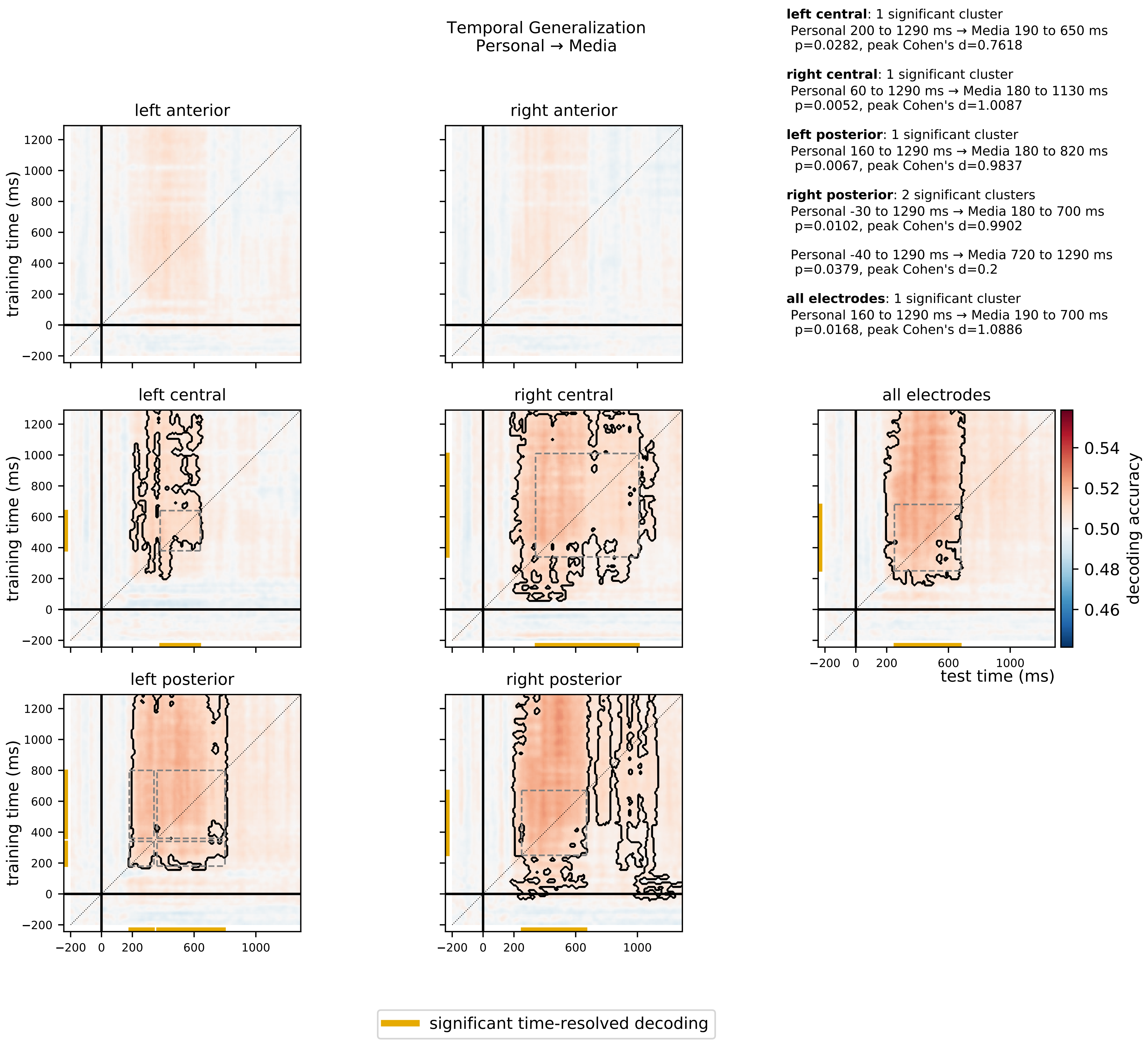

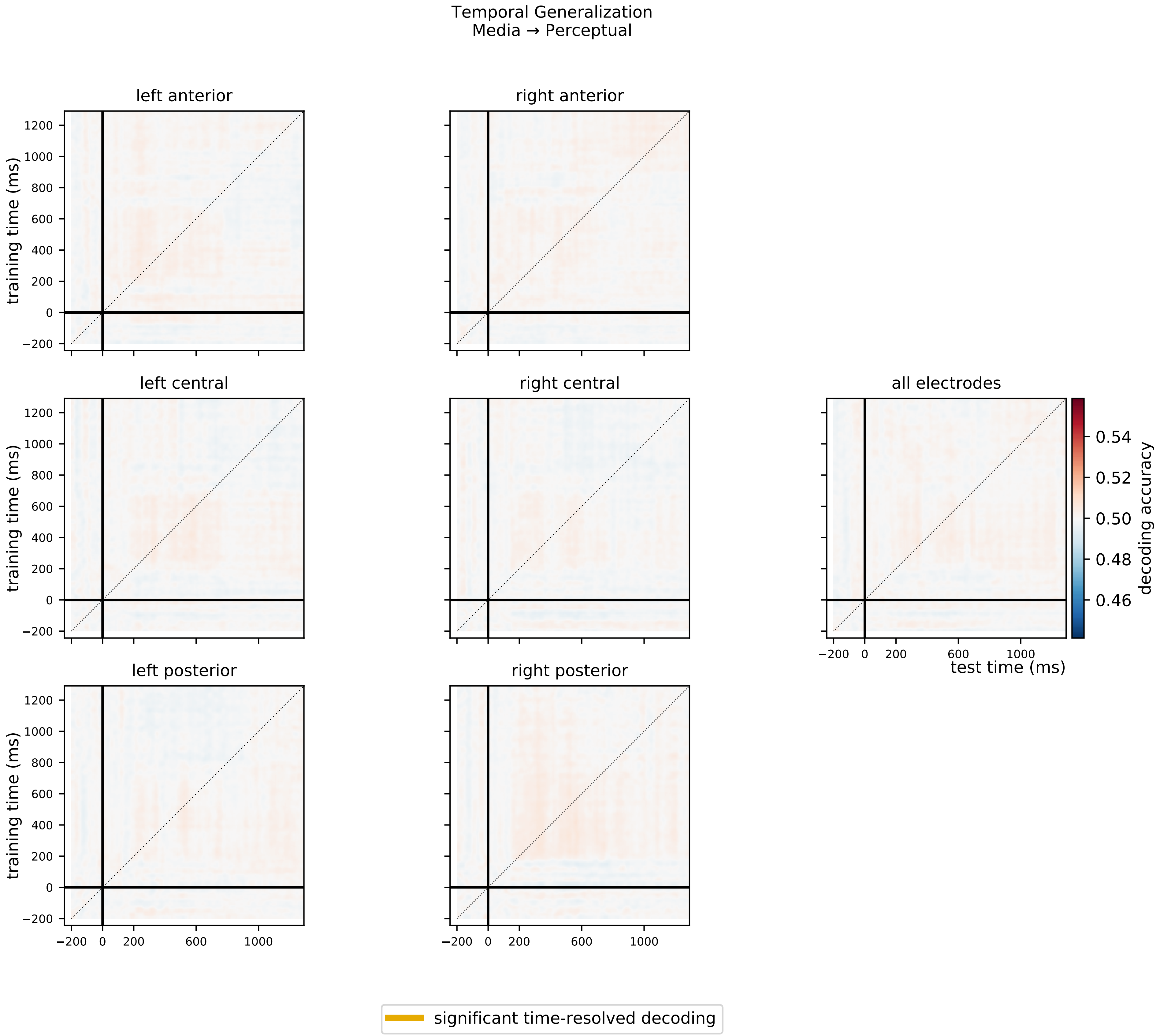

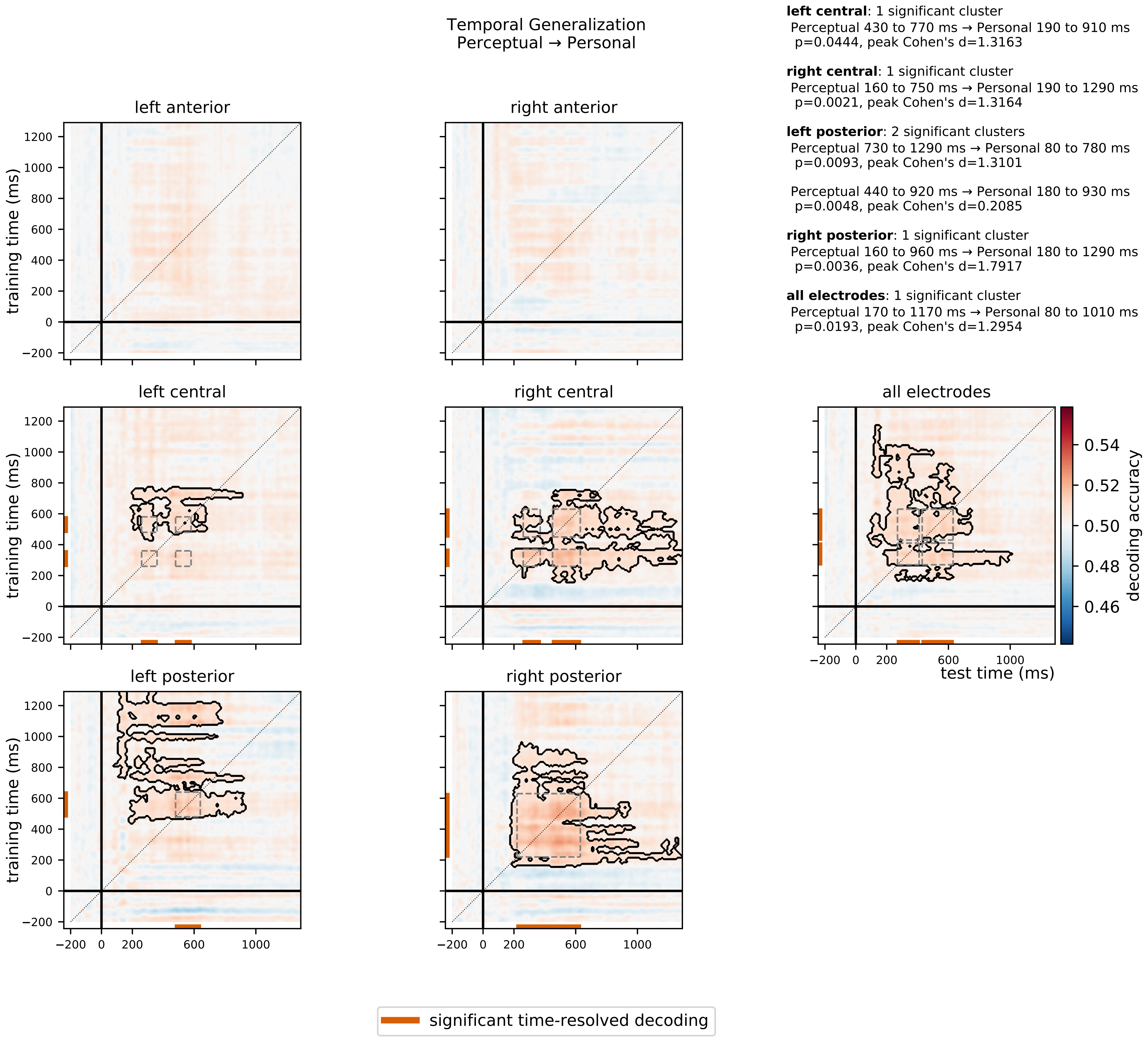

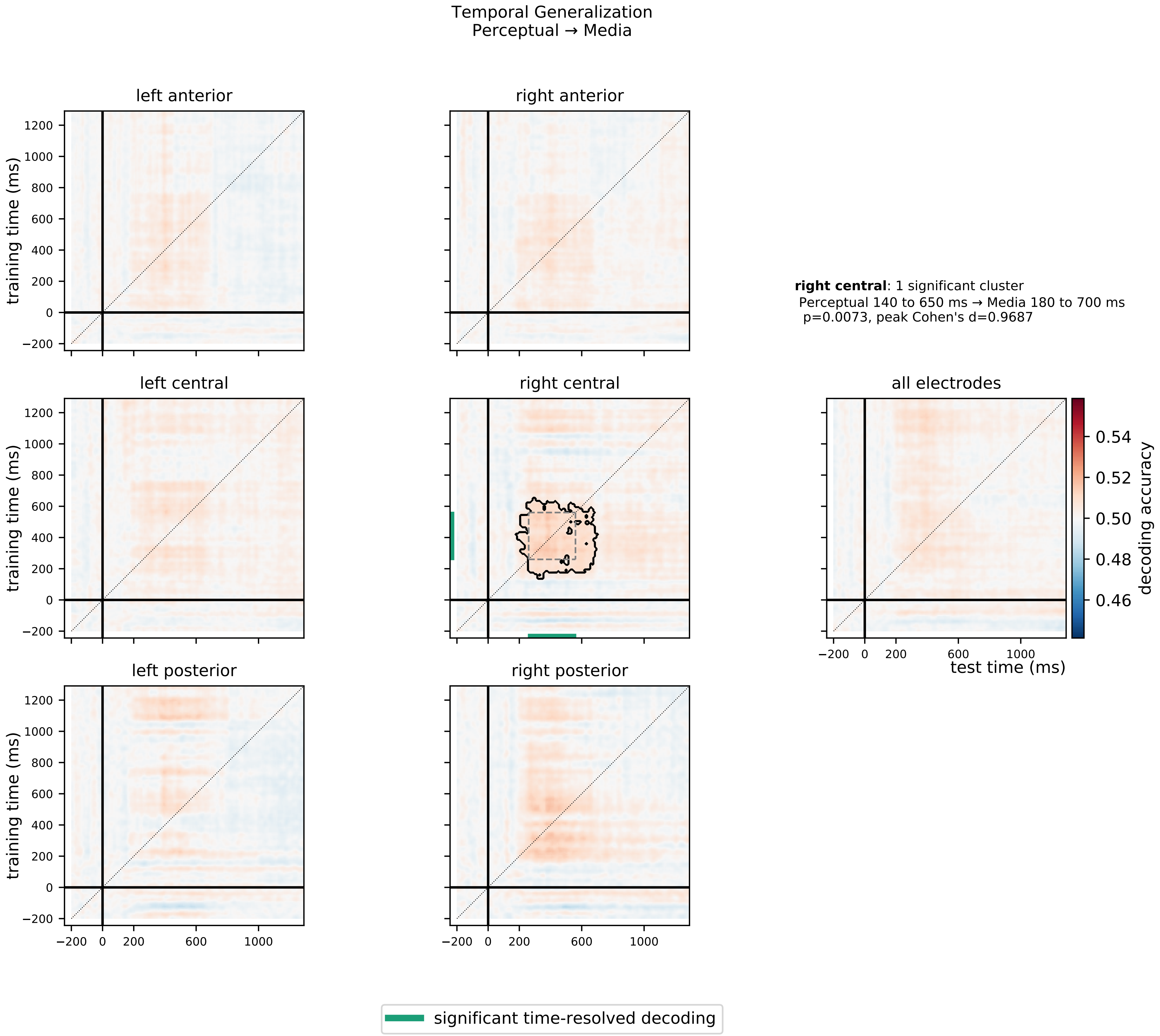

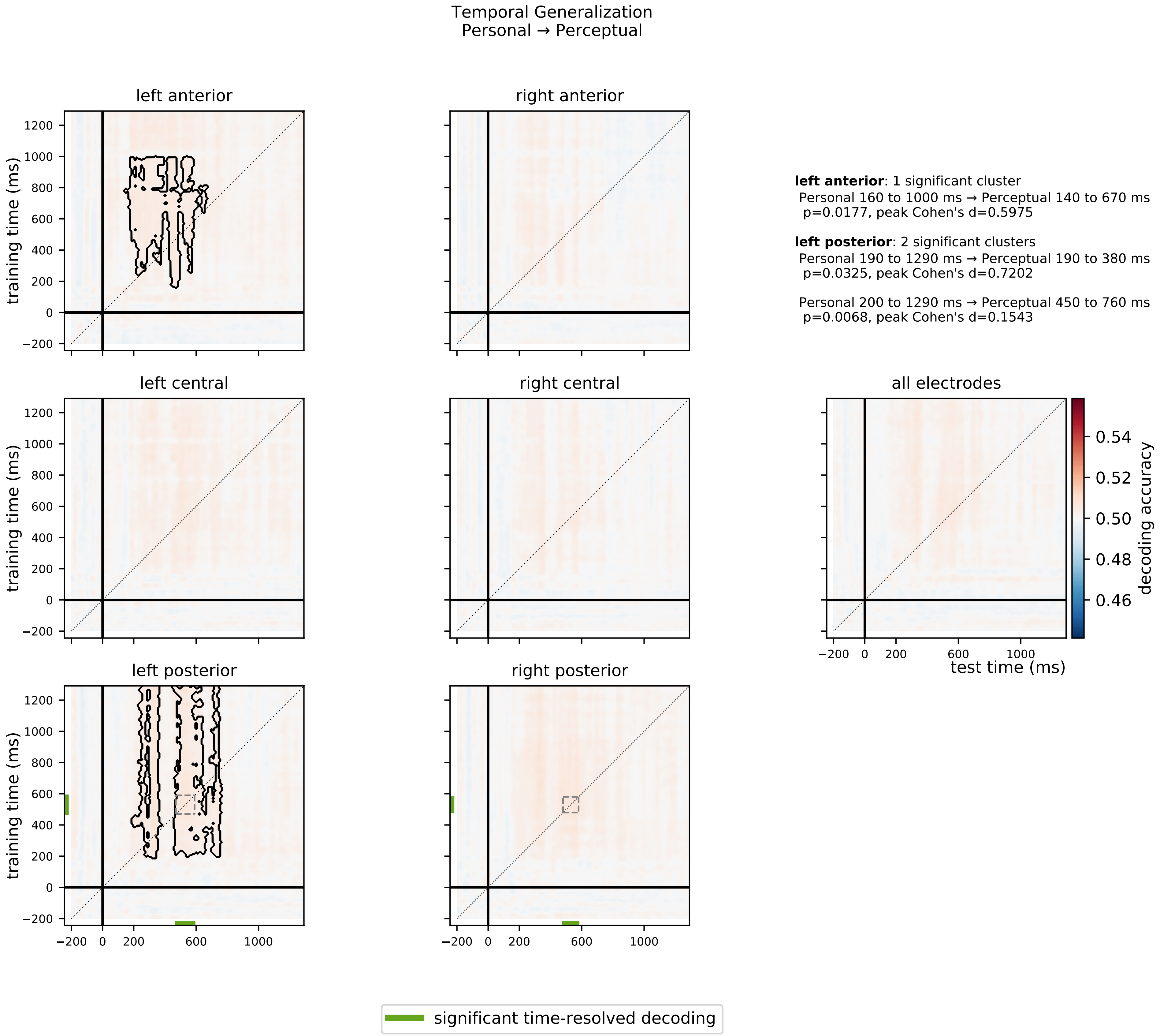

### Supplementary Figure 3

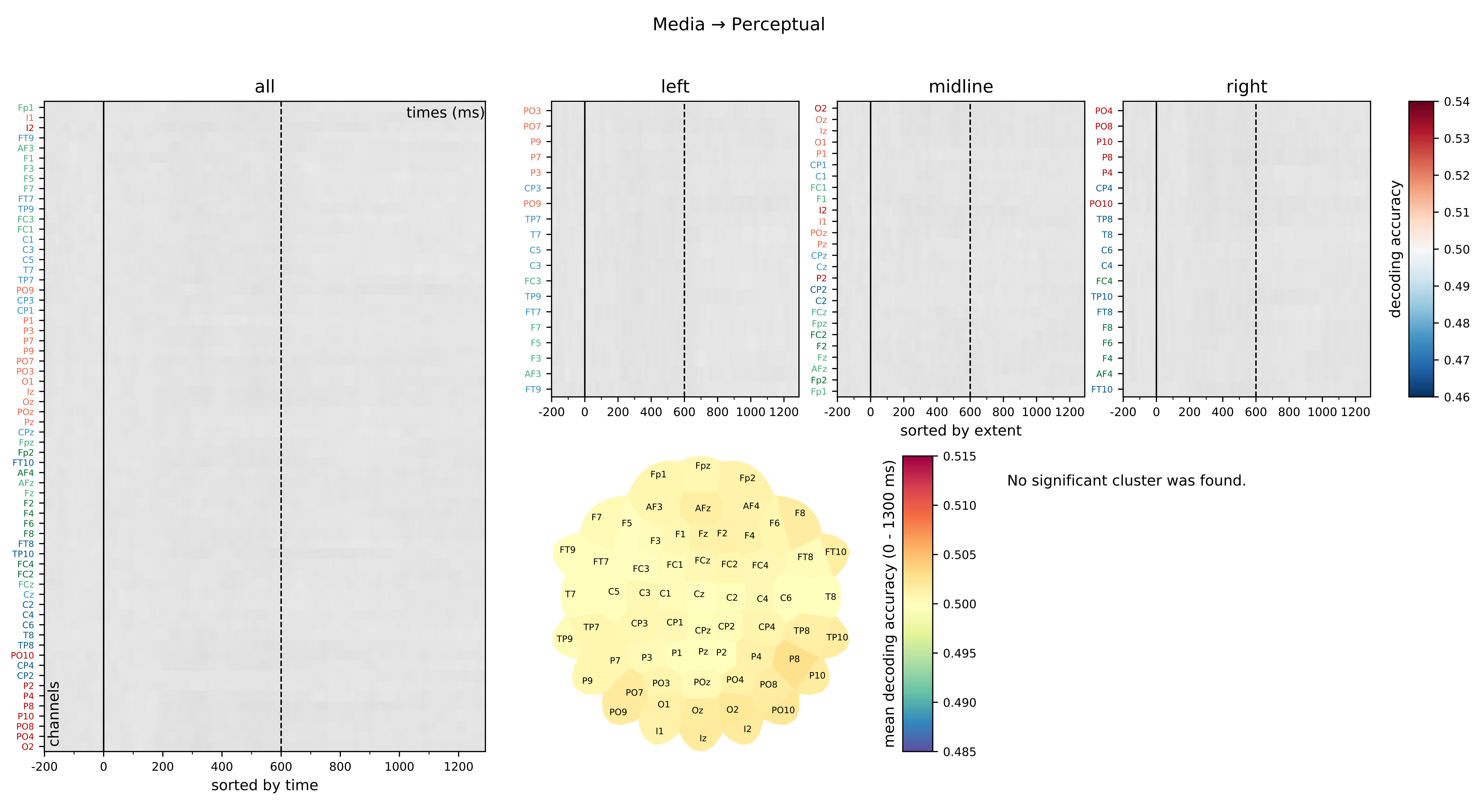

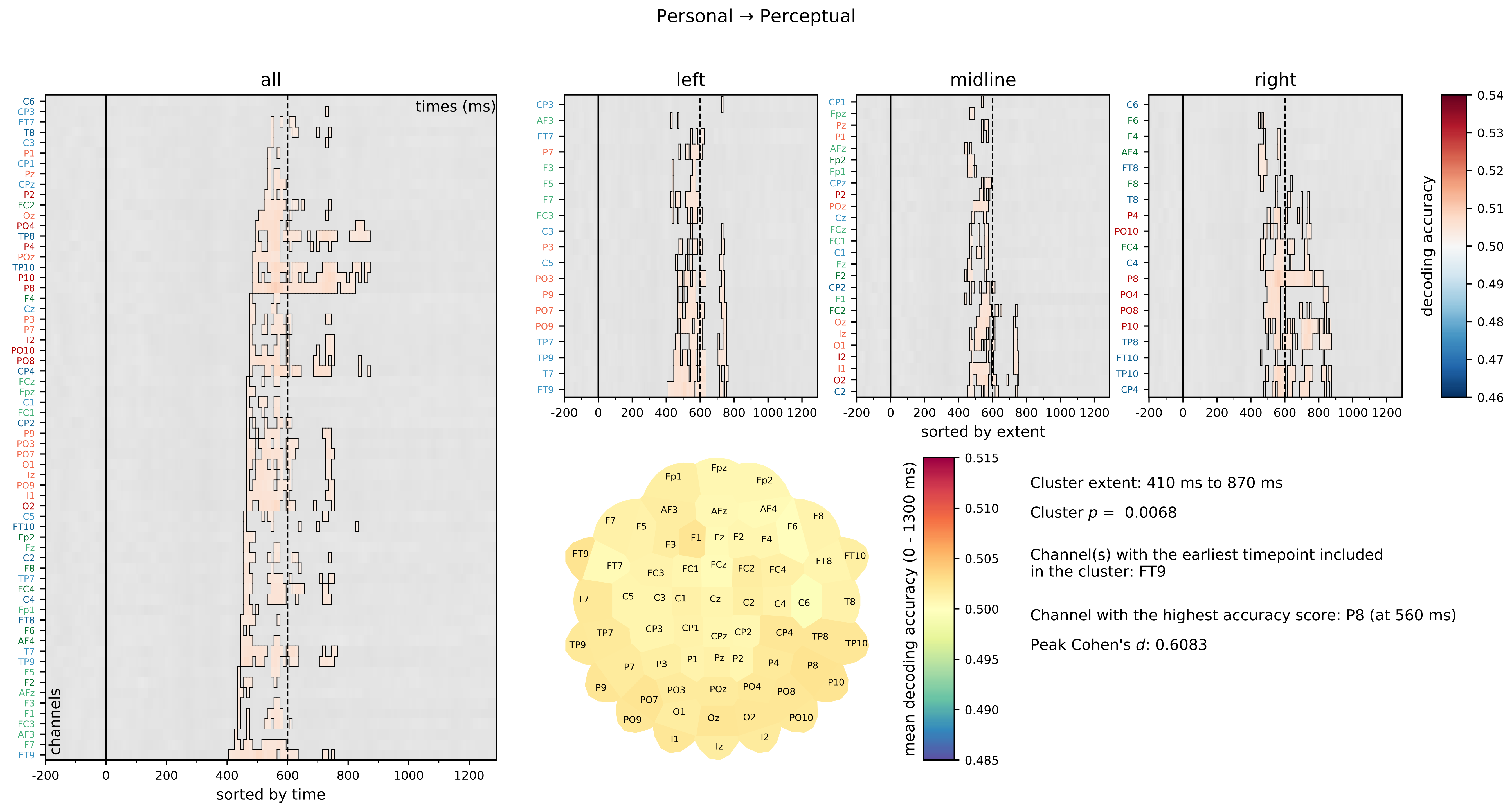

Perceptual → Media

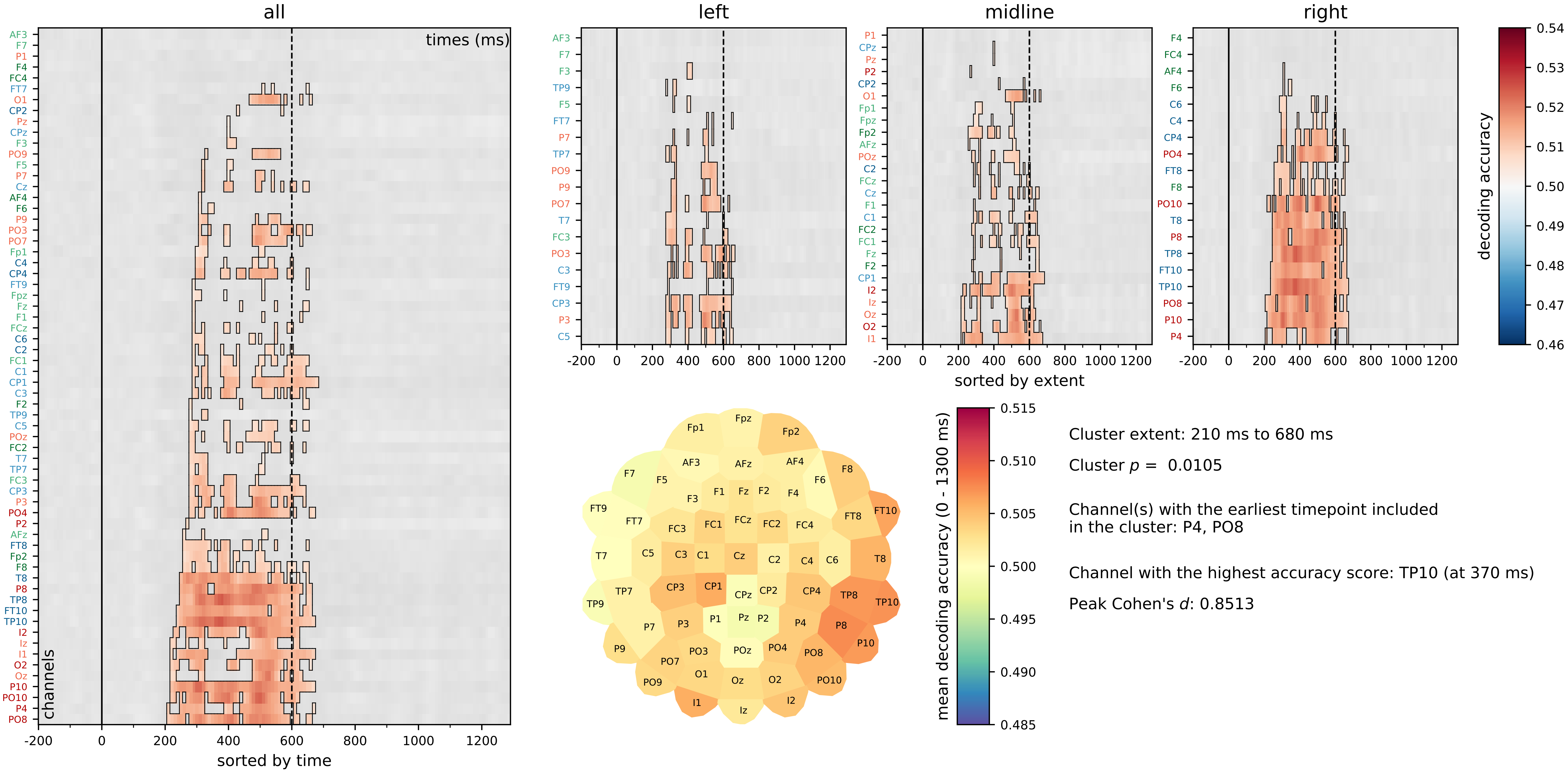

**Personal → Media**

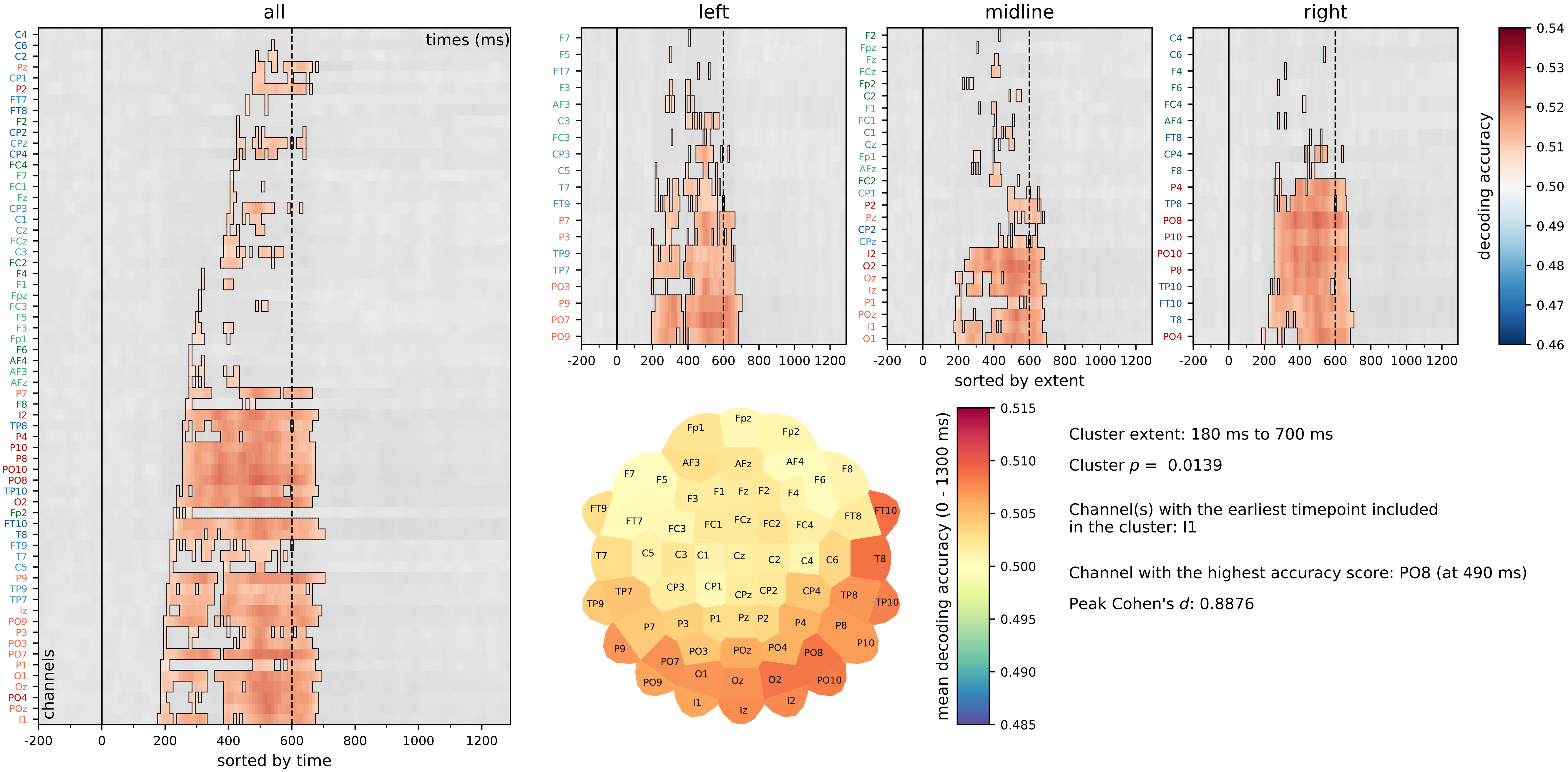

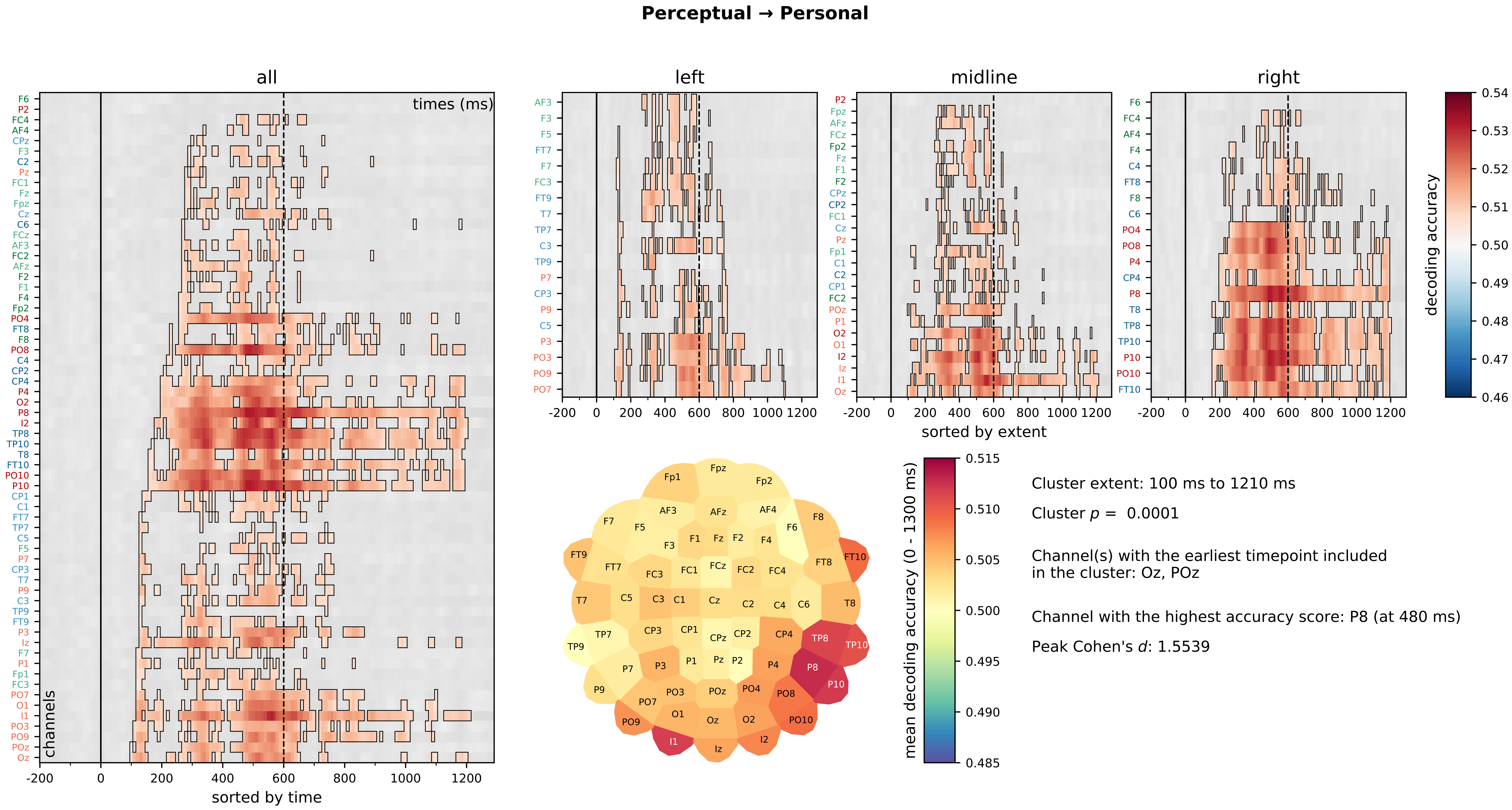

**Media → Personal**

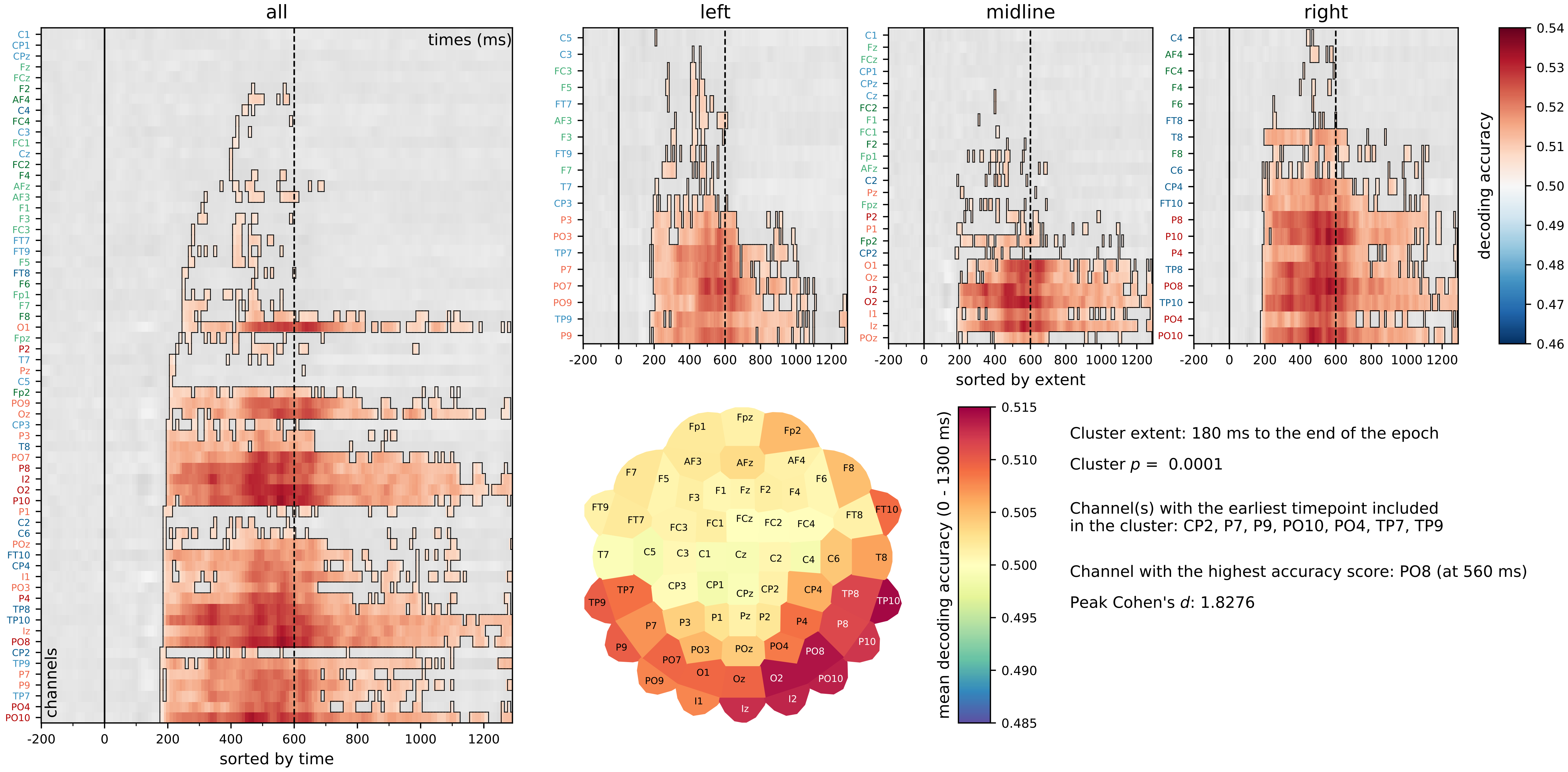
