## Supplementary Table 1 for "Evidence for a general neural signature of face familiarity"

### Difference from chance

**Perceptual→Media**

| | time window | peak latency | cluster *p* | peak Cohen's *d* |
|---|---|---|---|---|
| **all electrodes** | | | | |

| | left hemisphere | | | | right hemisphere | | | |
|---|---|---|---|---|---|---|---|---|
| | time window | peak latency | cluster *p* | peak Cohen's *d* | time window | peak latency | cluster *p* | peak Cohen's *d* |
| **anterior** | | | | | | | | |
| **central** | | | | | 260 - 560 ms | 320 ms | 0.0012 | 0.7186 |
| **posterior** | | | | | | | | |

**Perceptual→Personal**

| | time window | peak latency | cluster *p* | peak Cohen's *d* |
|---|---|---|---|---|
| **all electrodes** | 270 - 410 ms | 340 ms | 0.0264 | 1.0427 |
| | 430 - 630 ms | 470 ms | 0.011 | 0.9691 |

| | left hemisphere | | | | right hemisphere | | | |
|---|---|---|---|---|---|---|---|---|
| | time window | peak latency | cluster *p* | peak Cohen's *d* | time window | peak latency | cluster *p* | peak Cohen's *d* |
| **anterior** | | | | | | | | |
| **central** | 260 - 360 ms | 280 ms | 0.0316 | 0.7047 | 260 - 370 ms | 340 ms | 0.0065 | 0.9821 |
| | 480 - 580 ms | 500 ms | 0.0414 | 0.7241 | 450 - 630 ms | 550 ms | 0.0024 | 1.0644 |
| **posterior** | 480 - 640 ms | 520 ms | 0.0006 | 1.1029 | 220 - 630 ms | 490 ms | 0.0008 | 1.2744 |

**Media→Personal**

| | time window | peak latency | cluster *p* | peak Cohen's *d* |
|---|---|---|---|---|
| **all electrodes** | 190 - 1010 ms | 490 ms | 0.0004 | 1.5591 |

| | left hemisphere | | | | right hemisphere | | | |
|---|---|---|---|---|---|---|---|---|
| | time window | peak latency | cluster *p* | peak Cohen's *d* | time window | peak latency | cluster *p* | peak Cohen's *d* |
| **anterior** | | | | | | | | |
| **central** | | | | | 230 - 680 ms | 450 ms | 0.0006 | 0.8622 |
| **posterior** | 190 - 290 ms | 260 ms | 0.0447 | 0.8405 | 200 - 1290 ms | 470 ms | 0.0001 | 1.6381 |
| | 320 - 990 ms | 480 ms | 0.001 | 1.0333 | | | | |
| | 1010 - 1170 ms | 1100 ms | 0.0309 | 0.6807 | | | | |

**Personal→Perceptual**

| | time window | peak latency | cluster *p* | peak Cohen's *d* |
|---|---|---|---|---|
| **all electrodes** | | | | |

| | left hemisphere | | | | right hemisphere | | | |
|---|---|---|---|---|---|---|---|---|
| | time window | peak latency | cluster *p* | peak Cohen's *d* | time window | peak latency | cluster *p* | peak Cohen's *d* |
| **anterior** | | | | | | | | |
| **central** | | | | | | | | |
| **posterior** | 470 - 590 ms | 520 ms | 0.0148 | 0.4809 | 480 - 580 ms | 560 ms | 0.0427 | 0.5119 |

**Personal→Media**

| | time window | peak latency | cluster *p* | peak Cohen's *d* |
|---|---|---|---|---|
| **all electrodes** | 250 - 680 ms | 460 ms | 0.0112 | 0.6684 |

| | left hemisphere | | | | right hemisphere | | | |
|---|---|---|---|---|---|---|---|---|
| | time window | peak latency | cluster *p* | peak Cohen's *d* | time window | peak latency | cluster *p* | peak Cohen's *d* |
| **anterior** | | | | | | | | |
| **central** | 380 - 640 ms | 500 ms | 0.015 | 0.5935 | 340 - 1010 ms | 520 ms | 0.0026 | 0.8061 |
| **posterior** | 180 - 340 ms | 270 ms | 0.0382 | 0.5653 | 250 - 670 ms | 500 ms | 0.0072 | 0.8143 |
| | 360 - 800 ms | 520 ms | 0.005 | 0.9149 | | | | |

### Difference between decoding pairs

**Perceptual→Media - Media→Perceptual**

| | time window | peak latency | cluster *p* | peak Cohen's *d* |
|---|---|---|---|---|
| **all electrodes** | | | | |

| | left hemisphere | | | | right hemisphere | | | |
|---|---|---|---|---|---|---|---|---|
| | time window | peak latency | cluster *p* | peak Cohen's *d* | time window | peak latency | cluster *p* | peak Cohen's *d* |
| **anterior** | | | | | | | | |
| **central** | | | | | 270 - 450 ms | 380 ms | 0.0018 | -1.1265 |
| **posterior** | | | | | | | | |

**Perceptual→Personal - Personal→Perceptual**

| | time window | peak latency | cluster *p* | peak Cohen's *d* |
|---|---|---|---|---|
| **all electrodes** | | | | |

| | left hemisphere | | | | right hemisphere | | | |
|---|---|---|---|---|---|---|---|---|
| | time window | peak latency | cluster *p* | peak Cohen's *d* | time window | peak latency | cluster *p* | peak Cohen's *d* |
| **anterior** | | | | | | | | |
| **central** | | | | | 280 - 360 ms | 330 ms | 0.0237 | -0.9832 |
| **posterior** | 500 - 600 ms | 590 ms | 0.0428 | -0.7296 | 280 - 600 ms | 490 ms | 0.001 | -1.0386 |

**Media→Perceptual - Media→Personal**

| | time window | peak latency | cluster *p* | peak Cohen's *d* |
|---|---|---|---|---|
| **all electrodes** | 190 - 920 ms | 490 ms | 0.0001 | 1.6759 |

| | left hemisphere | | | | right hemisphere | | | |
|---|---|---|---|---|---|---|---|---|
| | time window | peak latency | cluster *p* | peak Cohen's *d* | time window | peak latency | cluster *p* | peak Cohen's *d* |
| **anterior** | | | | | | | | |
| **central** | | | | | 330 - 580 ms | 450 ms | 0.0011 | 1.0366 |
| **posterior** | 240 - 990 ms | 630 ms | 0.0001 | 1.4552 | 210 - 1290 ms | 470 ms | 0.0001 | 1.5921 |
| | 1010 - 1150 ms | 1100 ms | 0.0239 | 0.7937 | | | | |

**Personal→Perceptual - Personal→Media**

| | time window | peak latency | cluster *p* | peak Cohen's *d* |
|---|---|---|---|---|
| **all electrodes** | | | | |

| | left hemisphere | | | | right hemisphere | | | |
|---|---|---|---|---|---|---|---|---|
| | time window | peak latency | cluster *p* | peak Cohen's *d* | time window | peak latency | cluster *p* | peak Cohen's *d* |
| **anterior** | | | | | | | | |
| **central** | | | | | 380 - 700 ms | 520 ms | 0.0082 | 0.8696 |
| | | | | | 760 - 1010 ms | 960 ms | 0.0107 | 1.0114 |
| **posterior** | 370 - 720 ms | 490 ms | 0.0043 | 0.8047 | 280 - 660 ms | 380 ms | 0.0059 | 0.7511 |
